# SPARC in cancer-associated fibroblasts is an independent poor prognostic factor in non-metastatic triple-negative breast cancer and exhibits pro-tumor activity

**DOI:** 10.1101/2021.11.03.467104

**Authors:** Lindsay B Alcaraz, Aude Mallavialle, Caroline Mollevi, Florence Boissière-Michot, Hanane Mansouri, Joelle Simony-Lafontaine, Valérie Laurent-Matha, Thierry Chardès, William Jacot, Andrei Turtoï, Pascal Roger, Séverine Guiu, Emmanuelle Liaudet-Coopman

## Abstract

**Purpose:** Triple-negative breast cancer (TNBC) is the most aggressive breast cancer subtype and lacks specific targeted therapeutics. The current mechanistic evidence from cell-based studies suggests that the matricellular protein SPARC has a tumor-promoting role in TNBC; however, data on the clinical relevance of SPARC expression/secretion by tumor and stromal cells in TNBC are limited.

**Experimental Design:** This study analyzed the prognostic value of tumor and stromal cell SPARC expression in a large series of 148 patients with non-metastatic TNBC and long follow-up (median: 5.4 years). Fibrosis, tumor-associated macrophage (TAM) infiltration, tumor-infiltrating lymphocyte (TIL) density, PD-L1 and PD-1 expression were assessed. Tumor and stromal cell SPARC expression was studied by immunofluorescence, western blotting, and meta-analysis of published single-cell mRNA sequencing data. The biological role of fibroblast-secreted SPARC was analyzed using cell adhesion, wound healing, Transwell-based motility and invasion, and tumor spheroid assays.

**Results:** SPARC expression was detected in cancer cells (42.4%), cancer-associated fibroblasts (CAFs) (88.1%), TAMs (77.1%), endothelial cells (75.2%), and TILs (9.8%). Recurrence-free survival was significantly lower in patients with SPARC-expressing CAFs. SPARC expression in CAFs was an independent prognostic factor in multivariate analysis. Tumor and stromal cell SPARC expression was observed in TNBC cytosols, patient-derived xenografts, and cell lines. SPARC was expressed by different CAF subsets, including myofibroblasts and inflammatory CAFs. Fibroblast-secreted SPARC inhibited TNBC cell adhesion and stimulated their migration and invasion.

**Conclusions:** SPARC expression in CAFs is an independent predictor of recurrence-free survival in TNBC. Patients with SPARC-expressing CAFs could be eligible for anti-SPARC-targeted therapy.

**Statement of translational relevance:** Here, we identified a subgroup of patients with triple-negative breast cancer (TNBC) with worse prognosis and eligible for therapies that target extracellular matrix proteins in the tumor stroma. Specifically, we found that expression of the matricellular protein SPARC in cancer-associated fibroblasts (CAFs) is an independent prognostic marker of poor outcome in TNBC. Furthermore, we showed that in TNBC, SPARC is expressed by different CAF subpopulations, including myofibroblasts and inflammatory fibroblasts that are involved/associated with many tumor-related processes. We then found that SPARC secreted by fibroblasts has a pro-tumor-promoting role by inhibiting TNBC cell adhesion and stimulating their motility and invasiveness. Overall, our results support the need to consider SPARC expressed/secreted by CAFs as a novel therapeutic target in TNBC in the context of treatments to modulate the tumor stroma.

## Introduction

Triple-negative breast cancers (TNBC) are defined by the lack of estrogen receptor (ER), progesterone receptor (PR) and HER2 expression/amplification. TNBC represent 15% of all breast cancers [1]. Despite surgery, adjuvant chemotherapy and radiotherapy, TNBC prognosis is poor, mainly due to the disease heterogeneity and lack of specific therapeutic targets. TNBC is characterized by its unique tumor microenvironment that differs from that of other breast cancer subtypes and promotes cancer cell proliferation, angiogenesis, and drug resistance, while inhibiting apoptosis and tumor immune suppression [2]. TNBC microenvironment components, such as transformed extracellular matrix, soluble factors, immune cells, and re-programmed fibroblasts, hamper the host antitumor response and helps tumor progression and metastasis formation. In TNBC, stroma heterogeneity remains poorly understood, thus limiting the development of stromal cell-targeted therapies.

In the tumor microenvironment, heterogeneous populations of fibroblast-like cells, collectively termed cancer-associated fibroblasts (CAFs), are key players in the multicellular, stroma-dependent alterations that contribute to cancer initiation and progression [3]. Conversely, normal fibroblasts suppress tumor formation [4]. In breast cancer, CAF abundance has been associated with aggressive adenocarcinomas and predicts disease recurrence [5, 6]. In TNBC, recent single-cell RNA sequencing (scRNA-seq) studies highlighted a considerable CAF heterogeneity. Some CAF subpopulations have been characterized as key contributors to immune suppression, inflammation, and chemoresistance [7–10]. In breast cancer, tumor-associated macrophages (TAMs) are the most abundant inflammatory cells, and are typically M2-polarized cells with suppressive capacity [11] linked to their enzymatic activities and anti-inflammatory cytokine production [12]. TAMs support tumor progression and metastasis formation by blocking the anti-tumor immunity and by secreting factors that promote angiogenesis and epithelial-to-mesenchymal transition [11]. High M2-polarized TAM levels are associated with poorer TNBC outcome [13]. Tumor-infiltrating lymphocytes (TILs) constitutes a robust and independent prognostic marker in TNBC treated with (neo)adjuvant chemotherapy [14, 15]. TILs are associated with improved disease-free and overall survival (OS) rates in TNBC [16]. Programmed cell death (PD-1) (a CD-28-CTLA-4 family member) is an immune check-point receptor expressed by immune cells that contributes to the immune tolerance of self-antigens by peripheral T cells. PD-L1 (one of its ligand) is expressed by immune cells, epithelial breast cancer cells, and TILs. Activation of the PD-1-PD-L1 pathway specifically inhibits T-cell activation, and is one of the mechanisms that allow cancer cells to escape the antitumor immune response [17]. It is thought that TNBC are more immunogenic than other breast cancers. Indeed, the available evidence indicates that in TNBC, PD-L1 expression is more frequent (up to 60%) than in other breast cancers, and that PD-L1 tumor expression is positively associated with stromal TILs [18].

The matricellular protein Secreted Protein Acidic and Rich in Cysteine (SPARC; also known as osteonectin or basement membrane 40, BM40) is a Ca^2+^-binding glycoprotein that regulates extracellular matrix assembly and deposition, growth factor signaling, and cell-stroma interactions [19–22]. In cancer, SPARC is mainly secreted by neighboring stromal cells, but also by cancer cells [23–25]. SPARC plays oncogenic or tumor-suppressive roles, depending on the cancer type [26, 27]. In breast cancer, SPARC has pro-tumor functions and has been associated with worse prognosis [24, 28–33]; however, some studies also reported anti-tumor functions [34–36]. Mechanistic cell-based studies supports a tumor-promoting role in TNBC [37], suggesting that SPARC could be a candidate stromal therapeutic target.

The aim of this study was to evaluate SPARC expression in tumor and stromal cells, their prognostic value, and correlation with fibrosis, TAM infiltration, TIL density, PD-L1 and PD-1 levels in a large series of patients with non-metastatic TNBC. The objective was to identify a TNBC subgroup with worse prognosis and eligible for stroma-targeted therapy focused on extracellular matrix proteins.

## Materials and methods

### Antibodies and reagents

The rabbit polyclonal anti-SPARC (15274-1-AP) and the mouse monoclonal anti-periostin (clone No 1A11A3) antibodies were purchased from Proteintech. The mouse monoclonal anti-human SPARC (clone AON-5031, sc-73472) and the mouse monoclonal anti-HSC70 (clone B-6, sc-7298) antibodies were purchased from Santa Cruz Biotechnology. The mouse monoclonal anti-tubulin antibody (clone 236-10501, #A11126) was from Thermo Fisher Scientific. The horse anti-mouse immunoglobulin G (IgG)-horseradish peroxidase (#7076), and goat anti-rabbit IgG-HRP (#7074S) secondary antibodies were from Cell Signaling Technology. The donkey anti-goat HRP conjugated antibody (FT-1I7890) was from Interchim. The Alexa Fluor 488-conjugated anti-rabbit IgG (#Ab150077) was purchased from Abcam, and the Alexa Fluor 594-conjugated anti-mouse IgG (711-585-152) from ImmunoResearch Laboratories. Hoechst 33342 (#FP-BB1340) was from Interchim FluoProbes.

### Patients and tumor samples

For TNBC cytosols, patient samples were processed according to the French Public Health Code (law n°2004-800, articles L. 1243-4 and R. 1243-61), and the biological resources center has been authorized (authorization number: AC-2008-700; Val d’Aurelle, ICM, Montpellier) to deliver human samples for scientific research. TNBC tissue micro-arrays (TMAs) included tissue samples from 148 patients with unifocal, unilateral, non-metastatic TNBC who underwent surgery at Montpellier Cancer Institute between 2002 and 2012. TNBC samples were provided by the Biological Resource Center (Biobank number BB-0033-00059) after approval by the Montpellier Cancer Institute Institutional Review Board, following the French Ethics and Legal regulations for the patients’ information and consent. All patients were informed before surgery that their surgical specimens might be used for research purposes. Patients did not receive neoadjuvant chemotherapy before surgery. ER and PR negativity were defined as <10% expression by immunohistochemistry (IHC), and HER2 negativity was defined as IHC 0/1+ or 2+ and negative fluorescent/chromogenic hybridization *in situ*. This study was reviewed and approved by the Montpellier Cancer Institute Institutional Review Board (ID number ICM-CORT-2016-04). The study approval for patient-derived xenografts (PDXs) was previously published [38].

### Construction of TNBC TMAs

Tumor tissue blocks with enough material at gross inspection were selected from the Biological Resource Center. The presence of tumor tissue in sections was evaluated by a pathologist after hematoxylin-eosin (HE) staining of few sections. Two representative tumor areas were identified on each slide from which two malignant cores (1 mm in diameter) were extracted with a manual arraying instrument (Manual Tissue Arrayer 1, Beecher Instruments, Sun Prairie, WI, USA). After arraying completion, 4 μm sections were cut from the TMA blocks. One section was stained with HE and the others were used for IHC.

### TMA IHC

TMA sections were incubated with antibodies against SPARC (mouse monoclonal antibody; clone AON-5031, Santa Cruz Technology), cytokeratin 5/6 (mouse monoclonal, clone 6D5/16 B4, Dako), epidermal growth factor receptor (EGFR) (mouse monoclonal, clone 31G7, inVitroGen), PD-1 (mouse monoclonal, clone MRQ-22, BioSB), PD-L1 (rabbit monoclonal, clone SP142, Roche) and CD163 (mouse monoclonal, clone 10D6, BioSB) on a Autostainer Link48 platform (Dako) using the EnVision FLEX^®^ system (Dako) for signal amplification and diaminobenzidine tetrahydrochloride as chromogen. TMA sections were analyzed independently by two trained observers both blinded to the clinicopathological characteristics and patient outcomes. In case of disagreement, sections were revised by a third observer to reach a consensus. Results from duplicate cores, when available, were averaged. Basal-like phenotype was defined by cytokeratin 5/6 and/or EGFR expression (>10% of tumor cells). SPARC signal in cancer cells was scored as negative (<1% of stained cells), or positive (≥ 1% of stained cells). SPARC signal in CAFs, TAMs, endothelial cells, and TILs was scored as negative (<50% of stained cells), or positive (≥50% of stained cells). SPARC signal in normal epithelial breast tissue samples (N) was compared to the paired tumor sample (T) and scored as lower (N<T), equal (=), or higher (N≥T). TIL density (peritumoral and intratumoral) was evaluated on HE-stained sections, and was scored as: 0 (no TILs), 1 (rare TILs), 2 (moderate infiltrate, fewer TILs than tumor cells), and 3 (diffuse infiltrate, more TILs than tumor cells). Fibrosis was evaluated on HE-stained sections, and was scored as: 0 (no CAF), >20%, 20%-50%, >50% of fibrosis. PD-1 expression by TILs was scored as follows: not evaluable (no TILs), 0 (no stained TIL), 1 (<10% of stained TILs), 2 (10-50% of stained TILs) and 3 (>50% of stained TILs). PD-L1 expression in tumor cells was considered positive if detected in ≥1% of cells. TAM density was scored in CD163-stained sections and compared to the TIL density: 0 (no TAM), 1 (rare TAMs), 2 (moderate infiltrate, fewer TAMs than TILs), 3 (diffuse infiltrate, more TAMs than TILs).

### Immunofluorescence analysis

Paraffin-embedded patient-derived xenografts (PDX) tissue sections were deparaffined, rehydrated, rinsed, and saturated in PBS with 5% fetal calf serum (FCS) at 4 °C overnight. Sections were incubated with 1.2 μg/mL anti-SPARC rabbit polyclonal antibody (15274-1-AP) and 5 μg/mL anti-periostin mouse monoclonal antibody (1A11A3), followed by incubation with AlexaFluor 488-conjugated anti-rabbit IgG and AlexaFluor 594-conjugated anti-mouse IgG (1/400), respectively. Nuclei were stained with 0.5 μg/mL Hoechst 33342. Sections were imaged with a 63 X Plan-Apochromat objective on z stacks with a Zeiss Axio Imager light microscope equipped with Apotome to eliminate out-of-focus fluorescence.

### TNBC cytosols, cell lines, conditioned medium, and western blotting

TNBC cytosols were previously prepared and frozen [39]. The MDA-MB-453, MDA-MB-436, MDA-MB-468, Hs578T, BT-549 and HCC1806 TNBC cell lines were obtained from SIRIC Montpellier Cancer. The SUM159 TNBC cell line was from Asterand (Bioscience, UK). The MDA-MB-231 TNBC cell line was previously described [40]. Human mammary fibroblasts (HMFs) were provided by J. Loncarek and J. Piette (CRCL Val d’Aurelle-Paul Lamarque, Montpellier, France) [41], THP1 monocytes by L. Gros (IRCM, Montpellier), and human umbilical vein endothelial cells (HUVECs) by M. Villalba (IRMB, Montpellier). Cell lines were cultured in DMEM with 10% FCS (EuroBio), except the SUM159 cell line (RPMI with 10% FCS) and the THP1 cell line (RPMI with 10% decomplemented FCS, 10 mM HEPES, 1 mM sodium pyruvate and 50 μM β-mercaptoethanol). THP1 monocytes were differentiated into M0 macrophages by exposure to phorbol 12-myristate 13-acetate (100 ng/ml; Sigma Aldrich) for 48h. Then, cells became adherent and the medium was replaced with fresh medium supplemented with interleukin-4 (20 ng/ml) for 24h to induce differentiation of M0 macrophages to M2-polarized macrophages. For western blotting experiments, cell lysates were prepared in lysis buffer (50 mM HEPES [pH 7.5], 150 mM NaCl, 10% glycerol, 1% Triton X-100, 1.5 mM MgCl2, 1 mM EGTA) containing cOmplete™ Protease Inhibitor Cocktail (Roche, Switzerland), and centrifuged at 13,000 x g for 10 min. The corresponding conditioned media were centrifuged at 500 x g for 5min. Proteins from whole cytosols (20 μg) or cell lysates (30 μg) and conditioned media (40 μl) were separated on 13.5% SDS-PAGE and analyzed by immunoblotting with the anti-SPARC (clone AON-5031) and anti-tubulin antibodies using standard techniques. To prepare conditioned medium, HMFs were grown to 90% confluence in DMEM complemented with 10% FCS. Following washes with phenol red- and serum-free medium to remove serum proteins, cells were incubated in DMEM buffered with 50 mM HEPES [pH 7.5] and without FCS for 24h. Medium was harvested, and centrifuged at 1000 rpm for 5min, followed or not by SPARC depletion. Briefly, HMF conditioned medium was incubated with 5 μg of monoclonal anti-human SPARC antibody (clone AON-5031, sc-73472) overnight, and pre-absorbed to protein G-agarose at 4°C. Then conditioned medium (SPARC-immunodepleted or not) was filtered using 0.22 μm filters to eliminate cell debris. Cleared HMF conditioned medium (HFM CM) was collected and added to MDA-MB-231 cells for *in vitro* functional assays. SPARC immunodepletion was confirmed by western blotting

### ScRNA-seq data meta-analysis

Previously published scRNA-seq data from five patients with TNBC were used [8]. Processed 10X Genomics (Pleasanton, CA, USA) data, obtained from the European Nucleotide Archive under the accession code PRJEB35405, were loaded in R (4.0) and processed using the Seurat 3.4 package and default parameters [42]. Individual cell populations were annotated as published in the original scRNA-seq study [8] with minor modifications when appropriate.

### Cell adhesion, migration, and invasion assays

MDA-MB-231 cell adhesion was assessed as previously described [37]. Briefly, 96-well plates were coated with fibronectin (10 μg/ml; sc-29011; Santa Cruz Biotechnology) at 4°C overnight, and saturated with 1% bovine serum albumin (BSA) in PBS. MDA-MB-231 cells were detached with HyQTase (HyClone), washed in DMEM without FCS, and 5 10^4^ cells were then plated and incubated in serum-free HMF CM (SPARC-immunodepleted or not) at 37°C for 30min. Non-adherent cells were removed by flotation on a dense Percoll solution containing 3.33% NaCl (1.10 g/l), and adherent cells were fixed (10% [vol/vol] glutaraldehyde) using the buoyancy method [43]. Cells were stained with 0.1% crystal violet, and absorbance was measured at 570 nm. For migration and invasion assays, 8-μm pore Transwell inserts (polyvinyl pyrrolidone-free polycarbonate filters) in 24-well plates (Corning Inc., Corning, NY, USA) were coated with 10 μg/ml fibronectin (500 ng) (migration assays) or Matrigel (100 μg, Corning) (invasion assays) at 4°C for 24h. MDA-MB-231 cells were plated (5 10^4^ cells/well) in serum-free HMF CM (SPARC-immunodepleted or not) on the coated insert in the upper chamber. In these different assays, DMEM supplemented with 10% FCS was used as chemoattractant in the bottom chamber. After 16h, non-migrating/non-invading cells on the apical side of each insert were scraped off with a cotton swab, and migration and invasion were analyzed with two methods: (1) migrating/invading cells were fixed in methanol, stained with 0.1% crystal violet for 30min, rinsed in water, and imaged with an optical microscope. Two images of the pre-set field per insert were captured (x100); (2) migrating/invading cells were incubated with 3-(4,5-dimethylthiazol-2-yl)-2,5-diphenyltetrazolium bromide (MTT; 5 mg/ml, 1/10 volume; Sigma-Aldrich) added to the culture medium at 37°C for 4h. Then, the culture medium/MTT solution was removed and centrifuged at 10,000 rpm for 5min. After centrifugation, cell pellets were suspended in DMSO. Concomitantly, 300 μl of DMSO was added to each well and thoroughly mixed for 5min. The optical density values of stained cells (cell pellet and corresponding well) were measured using a microplate reader at 570 nm.

### Wound healing assay by live cell imaging

Before each experiment, MDA-MB-231 cells were grown to confluence in 96-well plates in a standard CO_2_ incubator. The 96-pin IncuCyte^®^ WoundMaker was used to simultaneously create precise and reproducible wounds by gently removing cells from the confluent monolayer. After washing, serum-free HMF CM (SPARC-immunodepleted or not) was added, plates were placed in the IncuCyte device and cell monolayers were scanned every hour. Wound width, wound confluence, and relative wound density were calculated using user-informed algorithms that are part of the IncuCyte™ software package. These algorithms identify the wound region and provide visual representations of the segmentation parameters.

### Tumor spheroids

To generate tumor spheroids, 5 × 10^3^ MDA-MB-231 cells/well were seeded in 150 μl complete medium in ultra-low attachment 96-well plates (Corning^®^ 96-well Clear Round Bottom Ultra-Low Attachment Microplate, NY, USA). Plates were centrifuged at 1000 rpm for 10min, and 3 days later each spheroid was embedded in collagen gel that included 1X DMEM, penicillin and streptomycin, 2% of SPARC-immunodepleted FCS, 3.75g/l sodium bicarbonate, 20 mM Hepes, 1 mg/ml rat collagen I, and 1.5 mM NaOH (qsp 150 μl/well in H_2_O). After 30min at 37°C, serum-free HMF CM (SPARC-immunodepleted or not) was added on the spheroid-containing polymerized collagen gel. MDA-MB-231 cell invasion area was analyzed in representative images with ImageJ.

### Statistical analyses

Data were described using means, medians and ranges for continuous variables, and frequencies and percentages for categorical variables. Data were compared with the Kruskal-Wallis test (continuous variables) and the chi-square or Fisher’s exact test, if appropriate (categorical variables). All tests were two-sided, and p-values <0.05 were considered significant. The median follow-up was calculated using the reverse Kaplan-Meier method. Relapse-free survival (RFS) and OS were estimated using the Kaplan-Meier method and compared with the log-rank test. RFS was defined as the time between the date of the first histology analysis and the date of the first recurrence at any site. Surviving patients without recurrence and patients lost to follow-up were censored at the time of the last follow-up or last documented visit. OS was defined as the time between the date of the first histology analysis and the date of death from any cause. Multivariate analyses were performed using the Cox proportional hazard model. Hazard ratios (HR) were given with their 95% confidence interval (CI). All statistical analyses were performed with the STATA 13.0 software (StatCorp, College Station, TX).

## Results

### In TNBC, SPARC is expressed in stromal and tumor cells

To determine SPARC expression in TNBC (tumor and stroma), TMAs were generated using samples from 148 patients with TNBC (Table 1). Their median age was 61.5 years (range 30.2-98.6), and 68.2% of them received adjuvant chemotherapy. Most TNBC (52.7%) were pT2, and 60.8% pN0. Moreover, 85.5% of tumors were ductal carcinomas, 6.9% lobular carcinomas, and 7.6% other histological types; 11% of tumors were classified as Scarff-Bloom-Richardson grade 1-2. A basal-like phenotype was observed in 61.9% of samples, and 66.9% of tumors expressed PD-L1. In 51.7% of tumors, TAMs were more abundant than TILs, and >20% of fibrosis was observed in 74.4% of tumors. SPARC expression (>50% of stained cells) in CAFs, TAMs, endothelial cells, and TILs was detected in 88.1%, 77.1%, 75.2%, and 9.8% of TNBC samples, respectively (Fig. 1A-B, Table 1). SPARC staining in tumor cells (>1% stained tumor cells) was observed in 42.4% of TNBC samples (Fig. 1A, Table 1). In 80% of samples, SPARC expression was lower in the adjacent normal breast tissue than in the tumor tissue (Fig. 1A and 1C).

**Table 1.** Clinicopathological characteristics of the whole TNBC population and SPARC expression status in cancer and stromal cells.

| Clinical and tumor characteristics | Whole population<br>*N* = 148 | Clinical and tumor characteristics | Whole population<br>*N* = 148 |
| --- | --- | --- | --- |
| **Age (years), median [min-max]** |  | **SPARC expression in TAMs** |  |
| < 55 years | 61.5 [30.2-98.6] | Negative | 27 (22.9%) |
| ≥ 55 years | 51 (34.5%) | Positive | 91 (77.1%) |
|  | 97 (65.5%) | Missing | 30 |
| **Tumor size** |  | **SPARC expression in endothelial cells** |  |
| T1 | 52 (35.1%) | Negative | 27 (24.8%) |
| T2 | 78 (52.7%) | Positive | 82 (75.2%) |
| T3/T4 | 18 (12.2%) | Missing | 39 |
| **Nodal status** |  | **SPARC expression in TILs** |  |
| N- | 90 (60.8%) | Negative | 74 (90.2%) |
| N+ | 58 (39.2%) | Positive | 8 (9.8%) |
|  |  | Missing | 66 |
| **Histological grade (SBR)** |  | **TIL density** |  |
| 1-2 | 16 (11.0%) | [0-1] | 42 (29.6%) |
| 3 | 130 (89.0%) | >1 | 100 (70.4%) |
| Missing | 2 | Missing | 6 |
| **Histology** |  | **PDL-1 expression in tumor cells** |  |
| Ductal | 124 (85.5%) | < 1% | 45 (33.1%) |
| Lobular | 10 (6.9%) | ≥ 1% | 91 (66.9%) |
| Other | 11 (7.6%) | Missing | 12 |
| Missing | 3 |  |  |
| **Adjuvant chemotherapy** |  | **PDL-1 expression in TILs** |  |
| No | 47 (31.8%) | 0 | 20 (14.9%) |
| Yes | 101 (68.2%) | ]0-10[ | 32 (23.9%) |
|  |  | [10-50[ | 40 (29.9%) |
|  |  | ≥ 50 | 42 (31.3%) |
|  |  | Missing | 14 |
| **Basal-like phenotype** |  | **PD1 expression in TILs** |  |
| ≤10% | 56 (38.1%) | 0 | 18 (12.9%) |
| Basal | 91 (61.9%) | < 10 | 30 (21.3%) |
| Missing | 1 | [10-50[ | 74 (52.9%) |
|  |  | ≥ 50 | 18 (12.9%) |
|  |  | Missing | 8 |
| **SPARC expression in tumor cells** |  | **Fibrosis** |  |
| Negative | 76 (57.6%) | 0 | 4 (3.0%) |
| Positive | 56 (42.4%) | < 20% | 31 (22.6%) |
| Missing | 16 | 20%-50% | 27 (19.7%) |
|  |  | > 50% | 75 (54.7%) |
|  |  | Missing | 11 |
| **SPARC expression in CAFs** |  | **TAMs (inflammation)** |  |
| Negative | 15 (11.9%) | 0/1 | 25 (17.5%) |
| Positive | 111 (88.1%) | 2 | 44 (30.8%) |
| Missing | 22 | 3 | 74 (51.7%) |
|  |  | Missing | 5 |

SBR: Scarff-Bloom-Richardson; CAFs: cancer-associated fibroblasts; TAMs: tumor-associated macrophages; TILs: tumor-infiltrating lymphocytes

**Figure 1.**
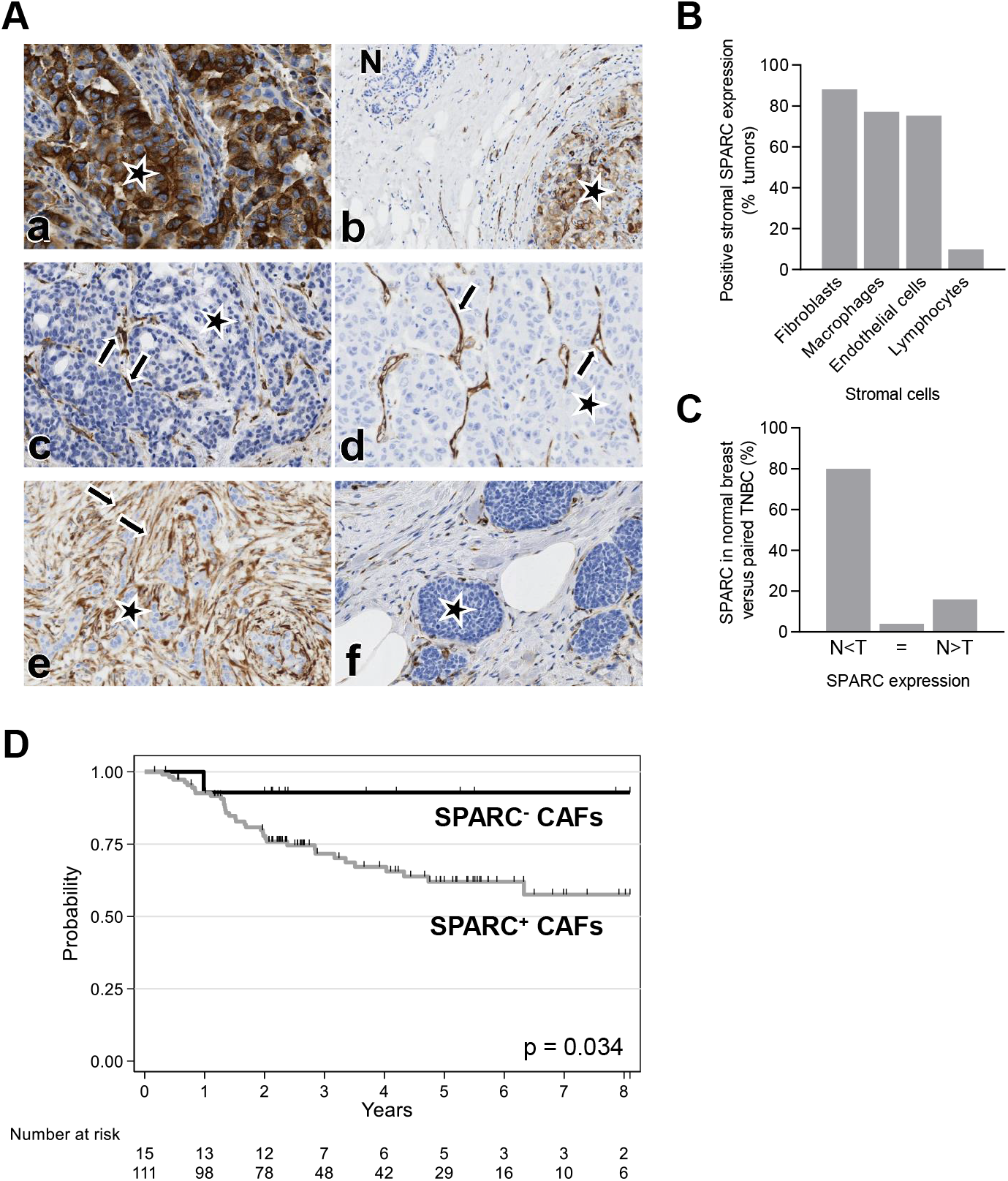
SPARC is a biomarker in TNBC and its expression in CAFs predicts RFS in TNBC. **(A) Representative images of TNBC tissue sections showing SPARC expression in cancer cells, CAFs, TAMs, endothelial cells, and in normal breast.** SPARC expression was analyzed in a TNBC TMA (n = 148 samples) by IHC using an anti-SPARC antibody (clone AON-5031). **(a)** SPARC expression in tumor cells. **(b)** Absence of SPARC expression in the adjacent normal breast tissue (N). **(c)** SPARC expression in TAMs. **(d)** SPARC expression in endothelial cells. **(e)** SPARC expression in CAFs. **(f)** Absence of SPARC expression in CAFs. SPARC scoring in cancer cells: positive (>1% of stained cells), negative (<1% of stained cells). SPARC scoring in stromal cells: positive (>50% of stained cells), negative (<50% of stained cells). Magnification X200. Stars: tumor cells; arrows: SPARC staining. **(B) Quantification of SPARC expression in TNBC stroma.** Percentage of TNBC samples with positive SPARC signal (>50% of stained cells) in the indicated stromal cell types. N = 148 samples. **(C) Quantification of SPARC expression in normal breast.** Percentage of normal breast tissue samples in which SPARC expression was lower (N<T), similar (=) or higher (N>T) than in the adjacent TNBC. T, tumor; N, normal breast; n = 50 samples. **(D) Relapse-free survival according to SPARC expression status in CAFs.** Patients with TNBC were divided in two subgroups according to SPARC expression in CAFs: SPARC^+^ CAFs and SPARC^-^ CAFs.

### SPARC expression in CAFs predicts RFS in patients with TNBC

As SPARC was expressed in the tumor and stromal compartments, its prognostic value was then evaluated. The median follow-up time was 5.4 years (range [0.1-14.3]). Local or regional recurrence occurred in 10 (7%) patients, and metastases (alone or with loco-regional recurrence) in 32 (22.5%) patients. RFS was not different in patients with SPARC-positive (SPARC^+^) and SPARC-negative (SPARC^-^) tumor cells (Supplementary Fig. S1) (Table 2). Conversely, RFS was lower in patients with SPARC^+^ than SPARC^-^ CAFs (3-year RFS rate: 72%, 95% CI [61.5-79.6] vs. 93% (95% CI [59.1-99.0]; p = 0.034) (Fig. 1D, Table 2). Moreover, RFS tended to be better in patients with SPARC^+^ than SPARC^-^ TAMs (3-year RFS rate: 81%, 95% CI [70.2-87.7]) vs. 62%, 95% CI [39.2-78.2]; p = 0.088) (Supplementary Fig. S2). SPARC expression status in endothelial cells (Supplementary Fig. S3) and TILs (Supplementary Fig. S4) did not have any prognostic value (Table 2). In univariate analysis, tumor size, nodal status, adjuvant chemotherapy, and SPARC expression in CAFs were correlated with RFS (Table 2). In multivariate analysis, only nodal status (HR = 2.96, 95% CI [1.48-5.94], p = 0.001), adjuvant chemotherapy (HR = 0.35, 95% CI [0.18-0.68], p = 0.002) and SPARC expression in CAFs (HR = 6.17, 95% CI [0.84-45.2], p = 0.015) were independent prognostic factors of RFS (Table 2). During the follow-up, 46 (31.1%) patients died among whom 11 (7.4%) without any TNBC recurrence. In univariate analysis, age (p = 0.027), tumor size (p <0.001), nodal status (p = 0.002), and adjuvant chemotherapy (p = 0.006) were associated with OS (Supplementary Table S1). In multivariate analysis, only tumor size (p = 0.05), nodal status (p = 0.008), and adjuvant chemotherapy (p <0.001) were independent prognostic factors of OS (Supplementary Table S1). Patients with SPARC^+^ CAFs (n =111, 88.1%) were younger (61.3% vs. 93.3%; p = 0.018) and tended to have ductal tumors (88.0% vs. 73.3%; p = 0.08) compared with patients with SPARC^-^ CAFs (Supplementary Table S2). In addition, SPARC^+^ TAMs and SPARC^+^ endothelial cells were detected more frequently in patients with SPARC^+^ than SPARC^-^ CAFs (80.6% vs. 41.7%, p = 0.007, and 78.0% vs. 50%, p = 0.026, respectively) (Supplementary Table S2). Fibrosis (>50%) was significantly less frequent in patients SPARC^+^ than SPARC^-^ CAFs (48.6% vs. 80%; p = 0.028) (Supplementary Table S2). PD-L1 expression (>50%) in TILs was more frequently detected in patients with SPARC^+^ than SPARC^-^ CAFs (34.8% vs. 15.4%; p = 0.049) (Supplementary Table S2). TIL density, PD-L1 expression in tumor cells and PD-1 expression in TILs were not significantly different between patients with SPARC^+^ and SPARC^-^ CAFs (Supplementary Table S2).

**Table 2.**
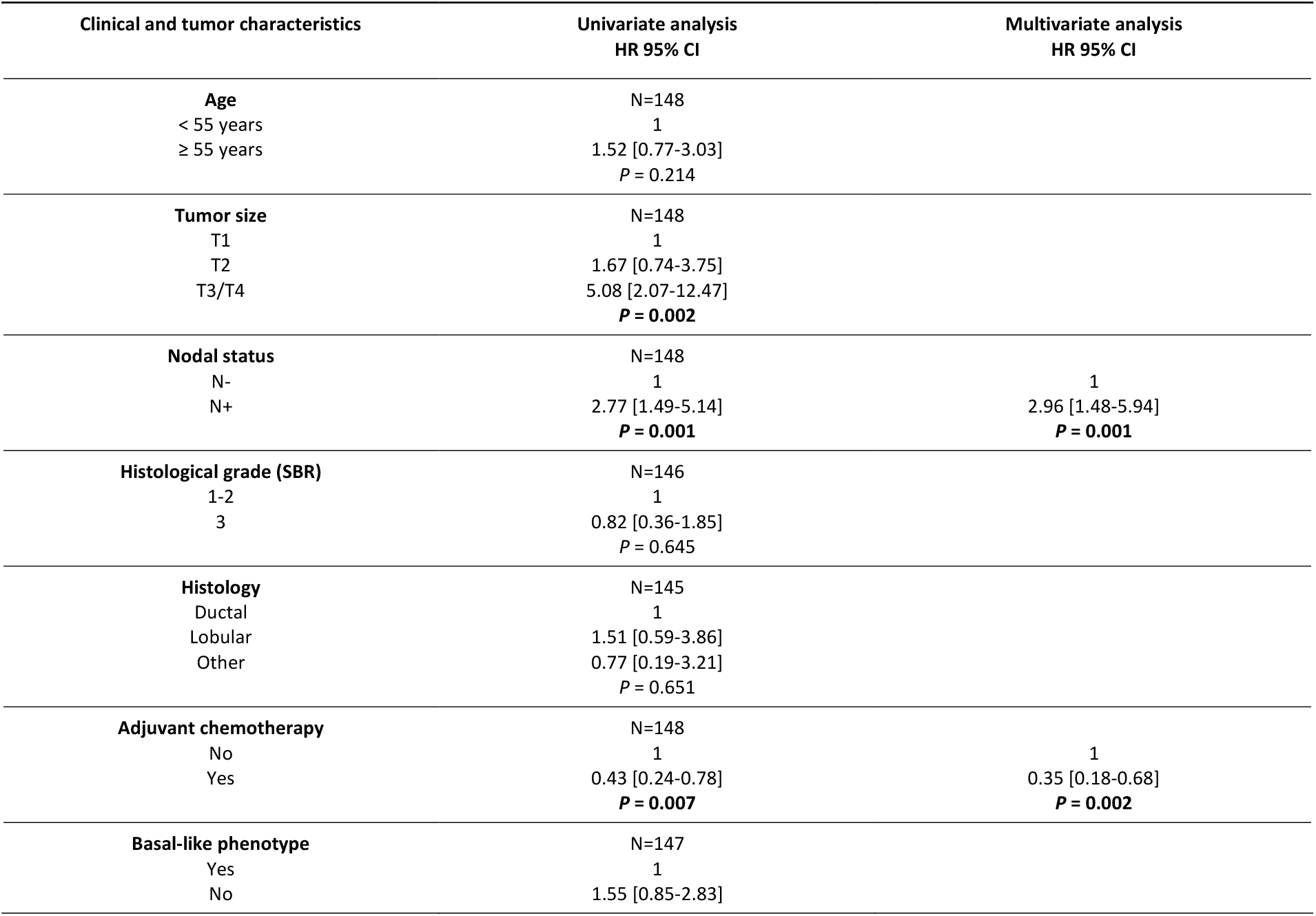
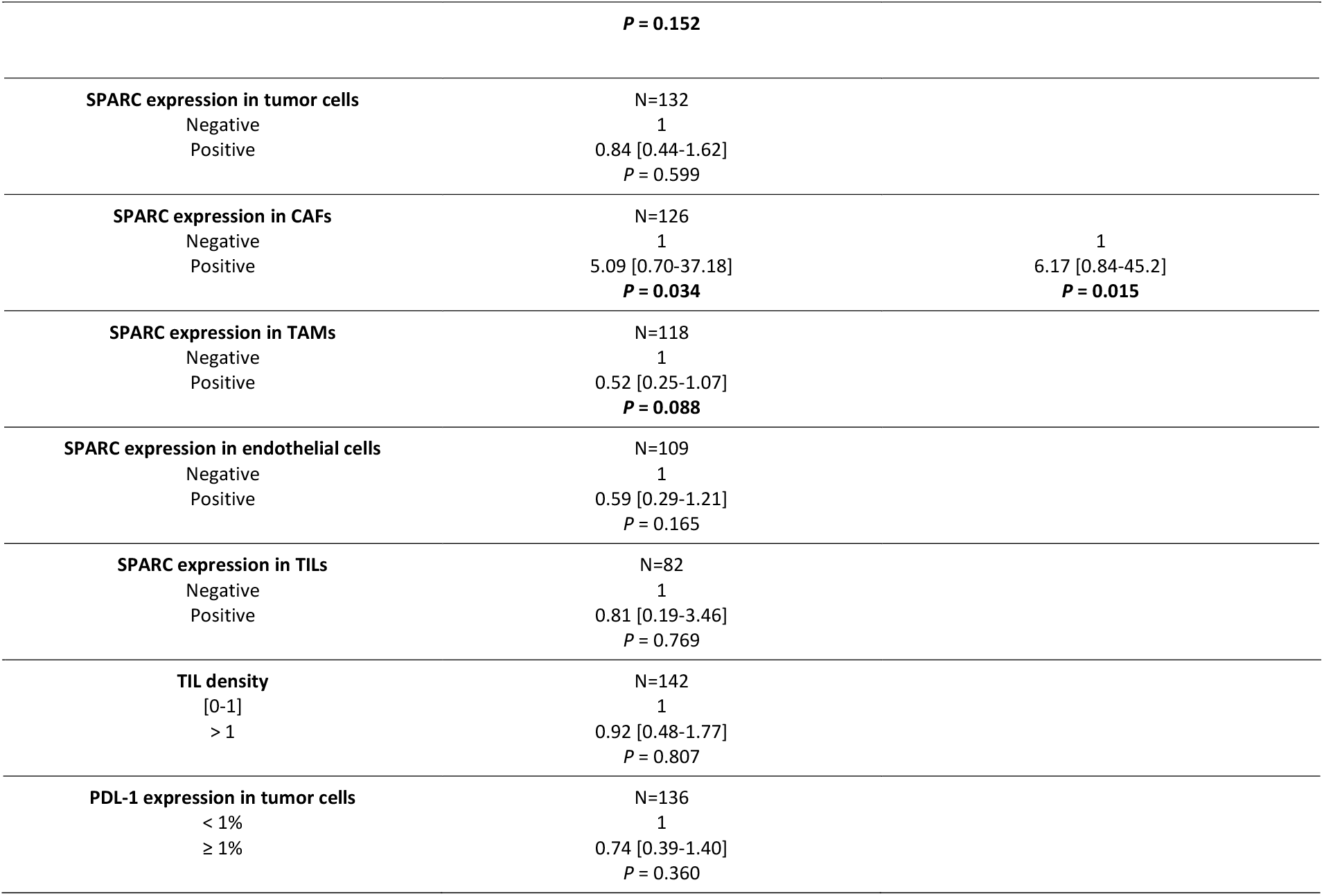
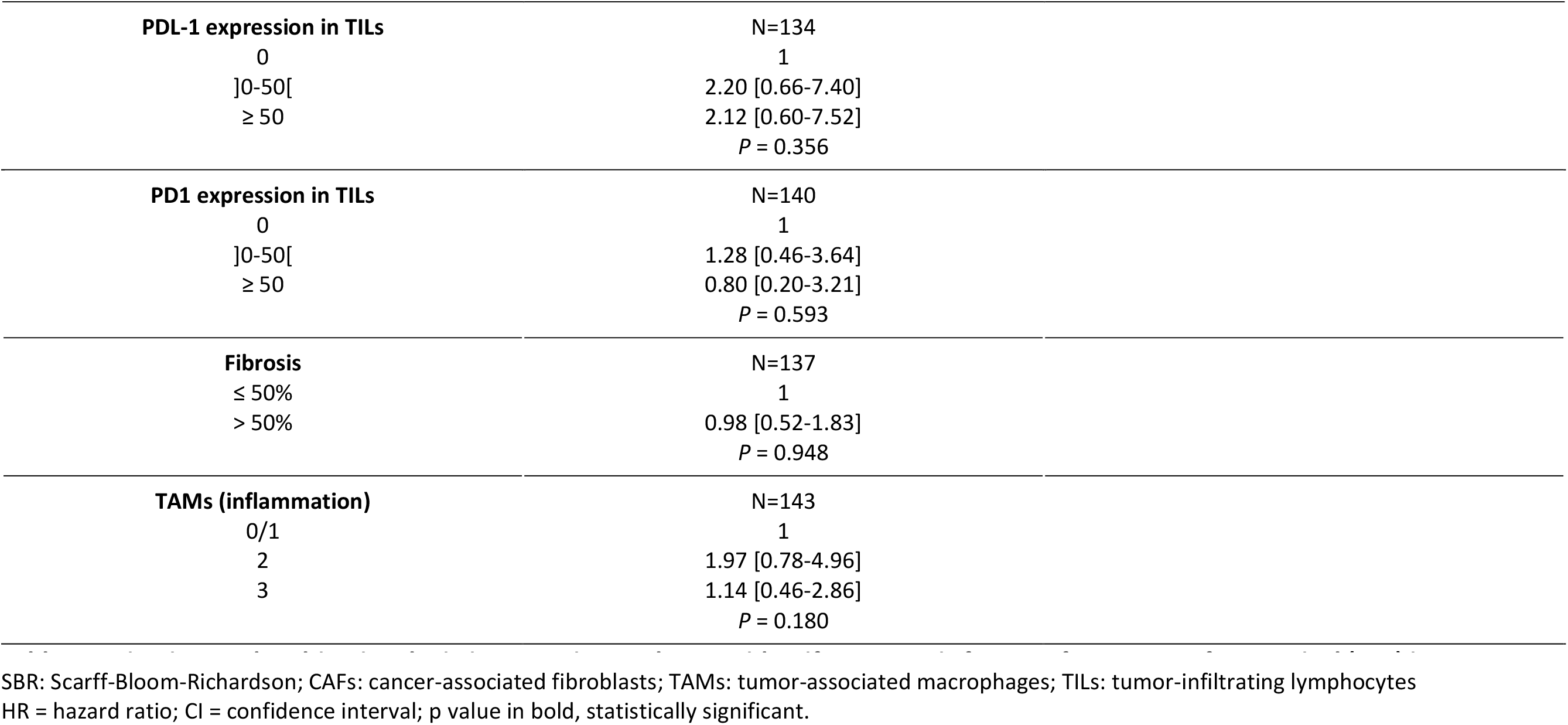
Univariate and multivariate logistic regression analyses to identify prognostic factors of recurrence-free survival (RFS) in TNBC.

### SPARC expression in TNBC cytosols, PDX, and cell lines

To further validate SPARC expression in TNBC, its expression was assessed in the cytosols of 30 primary TNBC samples by western blot analysis. SPARC protein was detected in all cytosols and SPARC cleaved fragments in about 30% of samples (Fig. 2A). SPARC protein expression and localization were then examined in two TNBC PDXs (PDX B1995 and PDX B3977) [38]. SPARC was localized in stromal cells, including CAFs, in the extracellular matrix, and in some tumor cells (Fig. 2B). Next, SPARC expression and secretion were analyzed in TNBC and stromal cell lines. SPARC was expressed and secreted by three of the eight TNBC cell lines tested (SUM159, Hs578T, BT-549) that exhibit a basal-like phenotype (Fig. 2C). SPARC was also expressed and secreted by HMFs, and to a lesser extent by HUVECs and M2-polarized THP1 macrophages (Fig. 2D and Supplementary Fig. S5)).

**Figure 2.**
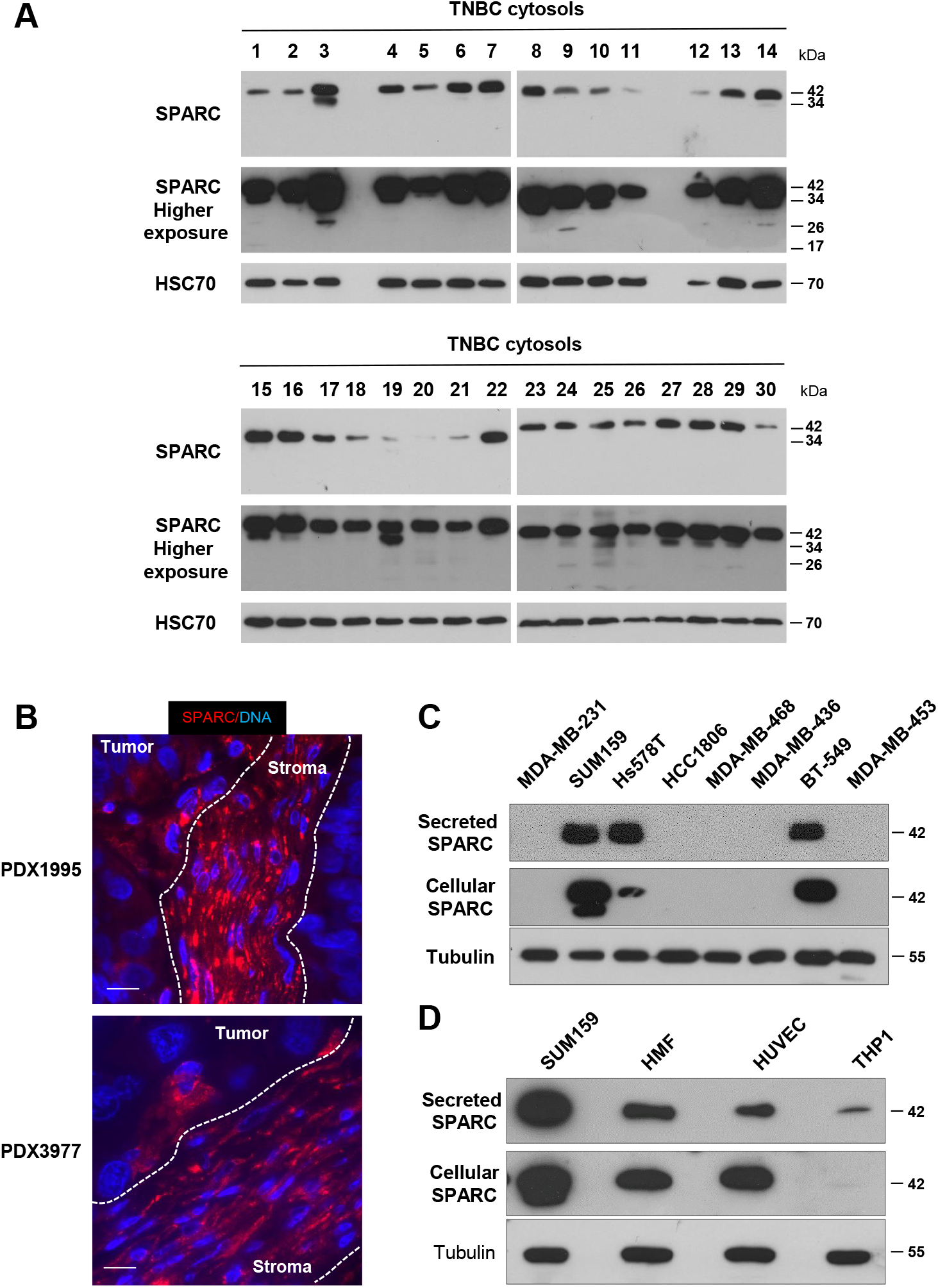
SPARC expression in TNBC cytosols, PDX, and cell lines. **(A) SPARC expression in TNBC cytosols.** SPARC expression was determined in 30 cytosols from primary TNBC biopsies. Whole cytosols (20 μg proteins) were analyzed by 13.5% SDS-PAGE and immunoblotting with an anti-SPARC antibody (clone AON-5031). A higher exposure of SPARC is shown. HSC70 (clone B-6) was used as loading control. **(B) SPARC expression and localization in TNBC PDX.** PDX B1995 and PDX B3977 sections were incubated with an anti-SPARC polyclonal antibody (15274-1-AP) (red). Nuclei were stained with Hoechst 33342 (blue). Scale bar, 10 μm. **(C) SPARC expression and secretion in TNBC cell lines** Whole cell extracts (30 μg proteins) and serum-free 24h conditioned media (40 μl) from the indicated TNBC cell lines were separated on 13.5% SDS-PAGE and analyzed by immunoblotting with an anti-SPARC (clone AON-5031) antibody. Tubulin was used as loading control. **(D) SPARC expression and secretion in stromal cell lines** Whole cell extracts (30 μg proteins) and serum-free 24h conditioned media (40 μl) from the indicated cell lines were separated on 13.5% SDS-PAGE and analyzed by immunoblotting with an anti-SPARC (clone AON-5031) antibody. Tubulin was used as loading control.

### SPARC is expressed in different CAF subsets

Based on the finding that SPARC expression in CAFs predicts RFS in TNBC, SPARC expression in different CAF subpopulations was thoroughly investigated through meta-analysis of recently published scRNA-seq data from patients with TNBC [8, 9]. In the first dataset (n= 5 patients with TNBC) [8], the t-distributed Stochastic neighbor embedding (tSNE) technique identified twenty different cell populations, including two fibroblastic cell populations, the first with features of myofibroblasts (myCAFs), and the second with an inflammatory phenotype (iCAFs) characterized by high expression of growth factors and immunomodulatory molecules (Fig. 3A). The scRNA-seq data analysis [8] showed that *SPARC* mRNA was strongly expressed in myCAFs and iCAFs, as well as *POSTN* (the gene encoding periostin, a CAF-secreted protein that promotes cancer progression and chemoresistance) [44] (Fig. 3B). *SPARC* was also detected in perivascular endothelial cells, myoepithelial cells, and basal cancer cells [8] (Fig. 3B), in accordance with our TMA analysis (Table 1). In the second scRNA-seq dataset (n=6 patients with TNBC) [9], high *SPARC* and *POSTN* mRNA levels were detected in three distinct CAF subtypes, in endothelial cells, M2-polarized macrophages, and cancer cells (where expression varied in function of the patient) (Supplementary Fig. S6), consistent with our TMA data (Table 1). As these two meta-analysis indicated that *SPARC* was expressed in different CAF subtypes, another scRNA-seq dataset (n=6 patients with breast cancer) that identified different myCAF and iCAF clusters was analyzed [10]. *SPARC* and *POSTN* mRNAs were detected mainly in myCAFs (ECM-myCAF, TGFβ-myCAF, Wound-myCAF, IFNαβ-myCAF, Acto-myCAF clusters), and also in iCAFs (IFNγ-iCAF, IL-iCAF, detox-iCAF clusters) (Supplementary Fig. S7). Altogether, these meta-analysis highlighted that *SPARC* mRNA is expressed by different CAF subtypes, including myofibroblasts and inflammatory-like CAFs involved in different tumor-related processes, such as matrix remodeling, inflammation, and resistance to therapy in TNBC [8, 10]. To complement the scRNA-seq findings, the localization of SPARC and periostin was investigated in the TNBC PDX B1995 microenvironment. Co-labeling with anti-SPARC and anti-periostin antibodies showed that SPARC (in red) partially co-localized with periostin (in green) in CAFs at the cancer cell-stromal interface (Supplementary Fig. S8).

**Figure 3.**
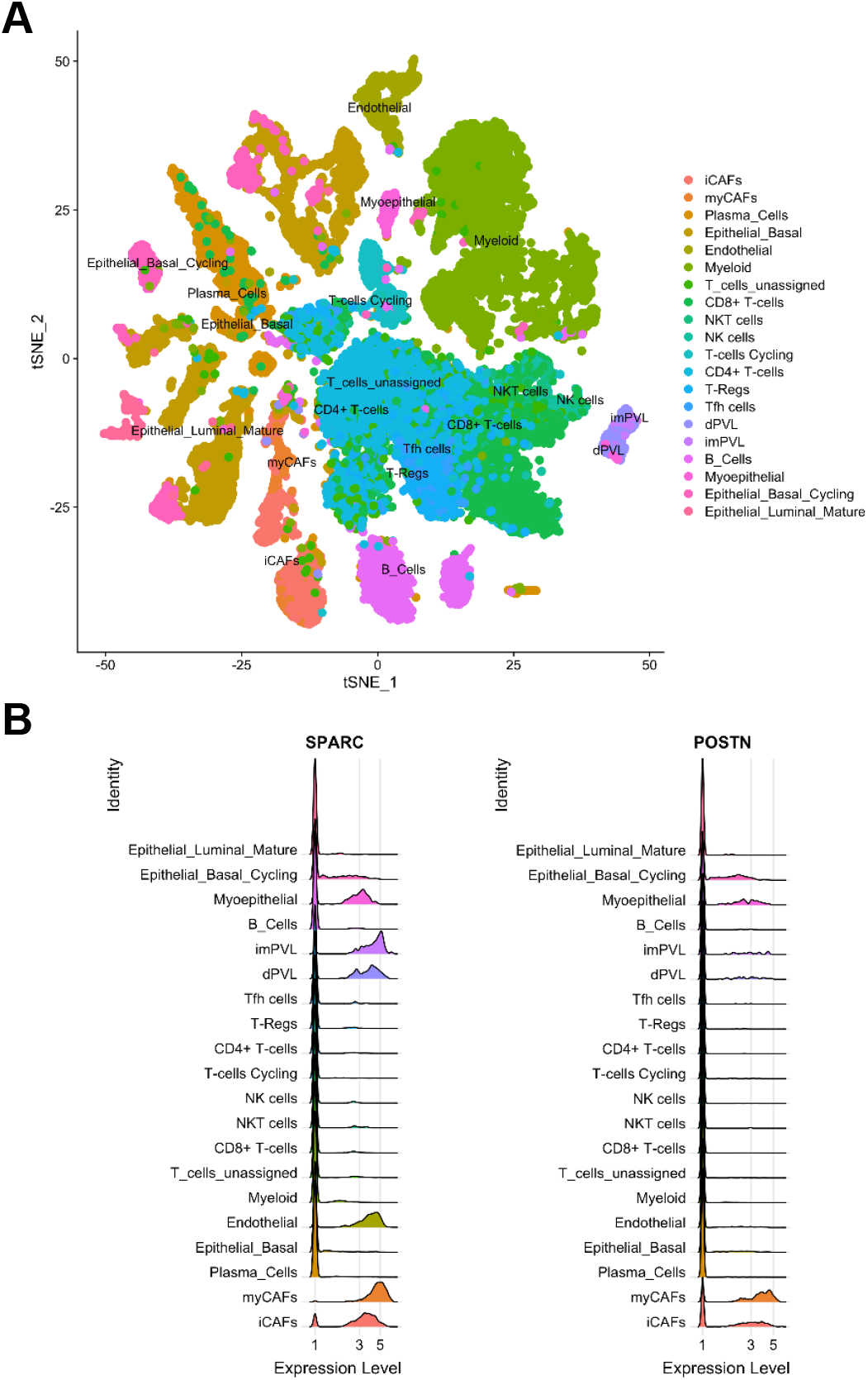
Expression of *SPARC* and *POSTN* mRNAs in TNBC by single-cell RNA-seq data analysis. **(A) Cell populations.** Twenty cell populations were identified by analysis of the previously published single-cell RNA-seq dataset PRJEB35405 that included five patients with TNBC, according to [8]. **(B) *SPARC* and *POSTN* mRNA expression.** Relative expression of *SPARC* and *POSTN* mRNA in each of the 20 populations identified by single-cell RNA-seq analysis, according to [8]. MyCAFs, myofibroblast-like CAFs; iCAFs, inflammatory-like CAFs; endothelial, endothelial cells; dPVL, differentiated perivascular-like cells; imPVL, immature perivascular-like cells; myoepithelial, myoepithelial cells; epithelial basal cycling, cancer cells.

### Fibroblast-secreted SPARC affects TNBC cell adhesion, migration and invasion

To obtain some insights into the pathophysiological relevance of SPARC^+^ CAFs in TNBC, the effects on TNBC cell adhesion, motility, wound healing, and invasiveness of SPARC-secreting HMF CM were investigated (Supplementary Fig. S9). The adhesion of MDA-MB-231 cells on fibronectin was reduced by 1.3-fold (*p <0.001*) after incubation with HMF CM compared with SPARC-immunodepleted HMF CM (Fig. 4A). Cell motility analysis in Boyden chambers showed that 88% of MDA-MB-231 cells passed through the fibronectin-coated filters after incubation with HMF CM (Fig. 4B). Motility was reduced by 2.3-fold when cells were incubated with SPARC-immunodepleted CM (Fig. 4B; *p <0.01*). Moreover, wound healing was significantly faster in MDA-MB-231 cells incubated with HMF CM than with SPARC-immunodepleted CM: wound closure was nearly complete after 16h in the presence of HMF CM (Fig. 4C). Lastly, MDA-MB-231 cell invasion through Matrigel-coated filters in Boyden chambers was 1.6-fold higher in the presence of HMF CM than SPARC-immunodepleted CM (Fig. 4D; *p <0.05*). The capacity of HMF-secreted SPARC to enhance MDA-MB-231 cell invasion was confirmed in a tumor spheroid assay (Fig. 4E). MDA-MB-231 tumor spheroid invasiveness at day 3 was 3.4-fold higher in the presence of HMF CM than SPARC-immunodepleted CM (Fig. 4E; *p <0.01*). Thus, HMF-secreted SPARC inhibits adhesion and promotes motility, wound healing and invasion of MDA-MB-231 TNBC cells, highlighting its pro-tumor role.

**Figure 4.**
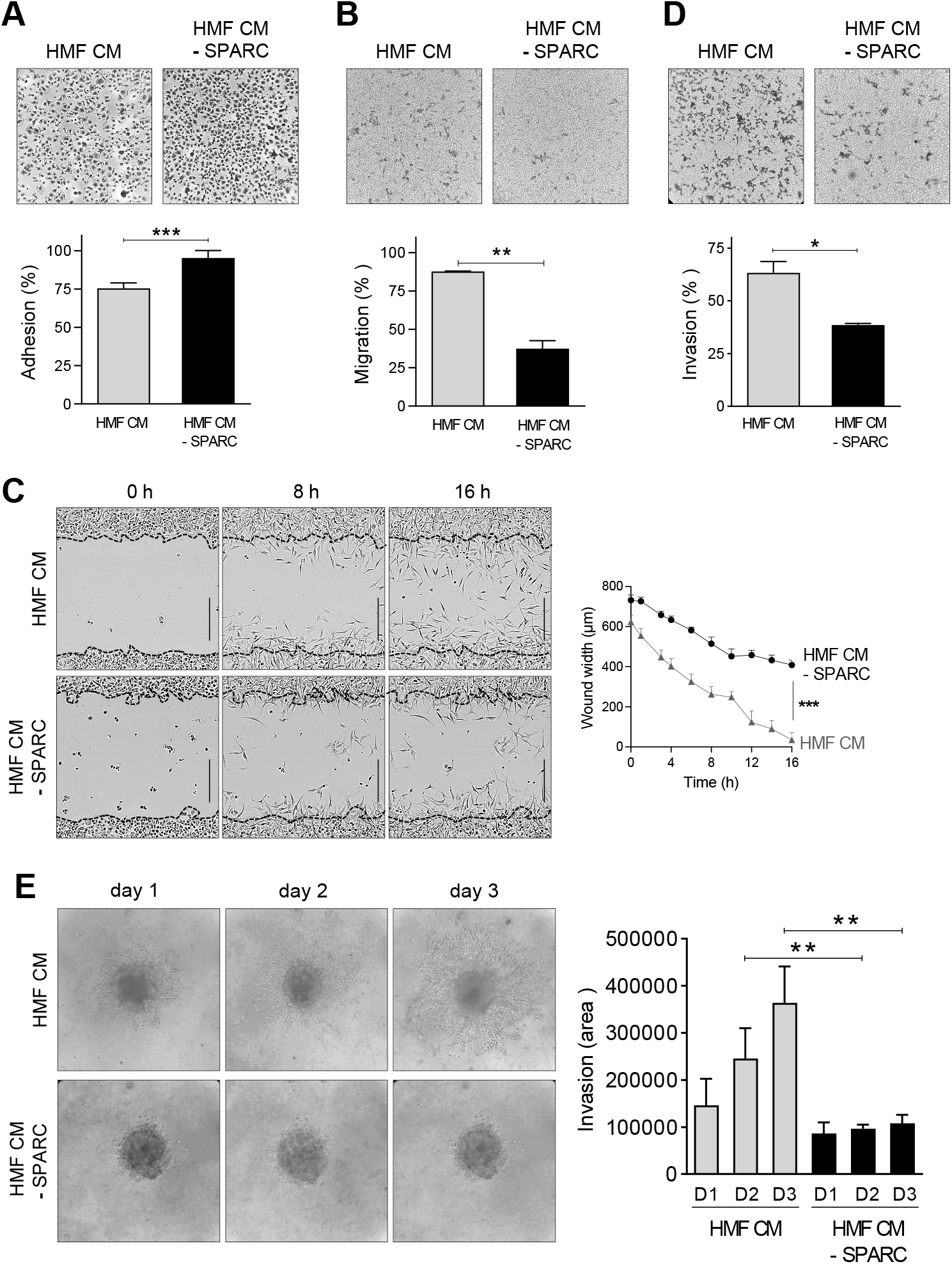
Effects of fibroblast-secreted SPARC on TNBC cell adhesion, migration and invasion. **(A) Cell adhesion.** MDA-MB-231 cells were let to adhere on a fibronectin matrix in the presence of CM HMF or SPARC-immunodepleted CM HMF (CM HMF IP SPARC) for 30 min. Upper panels, representative images of adherent cells stained with crystal violet. Lower panel, adhesion was quantified at 570 nm. Data are the mean (% of seeded cells) ± SD (n = 5); ***p <0.001 (Student’s *t*-test). Similar results were obtained in three independent experiments. **(B) Cell migration.** MDA-MB-231 cells were let to migrate for 16h on a fibronectin matrix in the presence of HMF CM or SPARC-immunodepleted HMF CM. Upper panels, representative images of migrating cells stained with crystal violet. Lower panels, quantification of migrating MTT-stained cells (absorbance was read at 570 nm). Data are the mean (% of seeded cells) ± SD (n = 3); **p <0.01 (Student’s *t*-test). Similar results were obtained in three independent experiments. **(C) Cell migration induced by wound healing.** MDA-MB-231 sub-confluent cell layers were wounded using the 96-well IncuCyte^®^ scratch wound assay. Left panels, representative images of MDA-MB-231 wound healing over time (t = 0 h, t = 6 h, t = 16 h) in the presence of HMF CM or SPARC-immunodepleted HMF CM. In the left panels, the initial scratch wound is delimited by the dashed lines. Bars, 400 μm. Right panel, wound healing (wound width, in μm) in the presence of HMF CM or SPARC-immunodepleted HMF CM was quantified over time. The data are the mean ± SD (n = 3); ***p <0.001 (Student’s *t*-test). Similar results were obtained in another independent experiment. **(D) Cell invasion.** MDA-MB-231 cells were let to invade on a Matrigel matrix in the presence of HMF CM or SPARC-immunodepleted HMF for 16h. Upper panels, representative images of invading cells stained with crystal violet. Lower panels, invading cells were stained with MTT and quantified at 570 nm. Data are the mean (% of seeded cells) ± SD (n = 3); ***p <0.001 (Student’s *t*-test). Similar results were obtained in three independent experiments. **(E) Cell invasion in tumor spheroid assay**. MDA-MB-231 tumor spheroids embedded in collagen I gel were let to invade in the presence of HMF CM or SPARC-immunodepleted HMF CM for 3 days. Left panels, representative images of invading MDA-MB-231 cells. Right panel, the invading MDA-MB-231 cell area was quantified using Image J. Data are the mean ± SD (n = 5); **p <0.01 (Student’s *t*-test).

## Discussion

Here, we showed that in TNBC, SPARC is expressed in both tumor and stromal cells, and that its expression in CAFs independently predicts RFS in patients with TNBC. Previous studies reported that SPARC is overexpressed in TNBC compared with other breast cancer molecular subtypes [45, 46]. In our study using IHC, SPARC expression in tumor cells was detected in 42% of TNBC samples, in agreement with previous literature data (SPARC expression in 37 to 52% of TNBC) [31, 45, 46]. However, SPARC expression in TNBC has never been correlated with clinicopathological parameters, such as age, histopathologic grade, tumor size and lymph node metastasis [31, 45]. Watkins *et al* reported that in breast cancer, SPARC is detected more frequently in ductal carcinomas [29]. Similarly, we found that ductal carcinoma tended to be more frequent in patients with SPARC^+^ CAFs, and that patients with TNBC with SPARC^+^ CAFs were often younger [47]. SPARC (mRNA or protein) overexpression prognostic value is controversial in TNBC. High SPARC expression in TNBC has been associated with poor prognosis in some studies [31, 33, 48], and with better prognosis in another [45]. We recently showed that high *SPARC* mRNA expression (n=225 patients with TNBC) tends to be associated with shorter RFS using an on line survival tool [37, 49]. In our current TNBC population, SPARC expression by tumor cells was not associated with RFS or OS. Studies using IHC reported that SPARC expression in tumor cells was associated with prognosis [31, 50]. Here, we found that SPARC was mainly expressed by stromal cells, including CAFs, and that its expression in CAFs was an independent prognostic factor of poor RFS in TNBC. In patients with SPARC^+^ CAFs, TILs more frequently expressed PD-L1, suggesting the interest to specifically evaluate the benefit of combining anti-PD1 or -PD-L1 with anti-SPARC targeted therapies in this TNBC subgroup. Moreover, fibrosis was less frequent in TNBC samples with SPARC^+^ CAFs, suggesting a better drug accessibility in this TNBC subgroup [51]. Other studies [47] reported a frequent SPARC stromal expression, but none, to our knowledge, evaluated its prognostic value or determined SPARC expression in the different stromal cell types.

Here, we observed the presence of SPARC cleaved fragments in about 30% of TNBC cytosols. The anti-SPARC antibody (clone AON-5031) used for IHC recognizes full-length SPARC and also some SPARC N-terminal fragments. Therefore, the prognostic value of SPARC expression in CAFs in TNBC described in the present study could be explained by the activity of the full-length protein and also of some of its cleaved fragments. SPARC includes three different structural and functional modules: the N-terminal acidic domain, the follistatin-like domain, and the C-terminal extracellular Ca^2+^ binding domain [21]. SPARC biological activity can be modulated by limited proteolysis, leading to the unmasking of distinct or amplified biological functions compared with those of the full-length protein [20, 52]. Matrix metalloproteinases (MMP-1, −2, −3, −9 and −13) cleave SPARC *in vitro* in its N-terminal acid domain and in its extracellular Ca^2+^ binding domain, releasing fragments that have higher affinity for collagens and that modulate cell-cell and cell-matrix extracellular interactions in the tumor microenvironment [53]. Moreover, MMP-3-mediated SPARC cleavage *in vitro* produces fragments that affect angiogenesis [54]. Cleavage of SPARC extracellular Ca^2+^ binding domain by MMP-8 and MMP-13 has been detected in the serum of patients with lung cancer, suggesting their presence also *in vivo* [55]. Similarly, cathepsin K cleaves SPARC *in vitro* and *in vivo* in its N-terminal acid domain and in its extracellular Ca^2+^ binding domain in mice harboring prostate cancer bone metastases [56]. We recently reported that secreted SPARC is cleaved by cathepsin D in TNBC, releasing a 9-kDa SPARC fragment with enhanced oncogenic properties [37].

The meta-analysis of previously published scRNA-seq datasets [7–10] showed that SPARC is expressed by different CAF subsets in TNBC. CAFs are the most abundant stromal cells in many cancers, including TNBC, and they are a phenotypically heterogeneous population, generally described as having a myofibroblastic phenotype (i.e. secretory and contractile cells that express α-SMA). Recently, it was found that fibroblast heterogeneity occurs in breast cancers and in TNBC [7–10]. Two myofibroblastic subsets (CAF-S1 and CAF-S4) differentially accumulate in TNBC [7]. CAF-S1 cells promote an immunosuppressive microenvironment [7], whereas CAF-S4 cells have pro-metastatic function [57]. More recently, a scRNA-seq approach in breast cancer identified eight clusters within the immunosuppressive CAF-S1 subset, subdivided in myofibroblasts-like and inflammatory-like CAFs [10]. Another scRNA-seq-based study identified myofibroblasts-like and inflammatory-like CAFs with immunomodulatory properties in TNBC [8]. By reanalyzing these scRNA-seq datasets [8–10], we noticed that *SPARC* mRNA was expressed by different CAF subsets, especially myofibroblasts-like and inflammatory-like CAFs, as well as *POSTN*, a gene encoding periostin, a protein that is secreted by CAFs with pro-tumor activity in breast cancer [44]. We then confirmed that SPARC and periostin (partially) co-localize in CAFs within the TNBC PDX microenvironment. Future studies will determine whether SPARC participates in the homeostasis of these different CAF subpopulations in TNBC, and whether SPARC has a different prognostic value when expressed in the different CAF subgroups in TNBC. In TNBC, CAFs regulate a number of tumor-promoting processes, including motility and invasion, drug resistance, inflammation and immunosuppression [7, 8, 57–59]. Our results showed that SPARC secreted by fibroblasts acts directly on TNBC cells by inhibiting their adhesion and promoting/facilitating their motility and invasiveness. This suggests that SPARC may be a therapeutic target in TNBC. Drugs that target CAFs have emerged as an important option for improving cancer therapies, and targeting CAF-derived extracellular matrix proteins has been proposed as an innovative anti-stromal therapy [60]. Our work strongly suggests that CAF-derived SPARC also may be a promising candidate for anti-stromal therapy.

### Conclusion

In this series, almost 88.1% of TNBC harbored SPARC^+^ CAFs and displayed distinct clinicopathological characteristics. SPARC expression in CAFs independently predicted worse RFS. This biomarker could be useful to identify a specific TNBC subgroup with worse prognosis. Furthermore, SPARC was expressed by different CAF subpopulations in TNBC, and fibroblast-secreted SPARC exhibited pro-tumor functions. Our results could have therapeutic implications for future anti-SPARC^+^ CAF targeted therapy.

## Acknowledgements

This work was supported by a public grant overseen by the French National Research Agency (ANR) as part of the “Investissements d’Avenir” program (reference: ANR-10-LABX-53-01), SIRIC Montpellier Cancer Grant INCa_Inserm_DGOS_12553, University of Montpellier, the associations ‘Ligue Régionale du Gard’, ‘Ligue Régionale de l’Herault’, ‘Ligue Régionale de la Charente Maritime’, and ‘Association pour la Recherche sur le Cancer’ (ARC). We thank Prof Marc Ychou (Director of ICM, Montpellier, France) for the ICM financial support for Dr Lindsay Alcaraz’ salary.

## Authors’ contributions

## Supplementary Material and Methods

### Single-cell RNA-seq meta-analysis

Previously published single-cell RNA-seq data (n=6 patients with TNBC) and deposited in the Gene Expression Omnibus database under accession code GSE118390 [9] were used. They were loaded in R (4.0) and then processed using the Seurat 3.4 package using default parameters [42]. Individual cell populations were annotated as published in the original single-cell RNA-seq study [9] except for cancer cells that were labeled Cancer_P1, _P2, _P3, _P4, _P5, _P6. Previously published single-cell RNA-seq data (n=6 patients with breast cancer) deposited at the European Genome-Phenome Archive (EGA) platform (https://ega-archive.org) under accession number AS00001004031 [10] also were analyzed. Individual CAF-S1 clusters were annotated as published in the original study [10].

### RT-qPCR

RNA was isolated using the RNeasy^®^ Plus Mini Kit (Qiagen, Hilden, Germany) and 1 μg of total RNA was reverse transcribed using the SuperScript™ III reverse transcriptase kit (Invitrogen). Real-time PCR was performed using SYBR^®^ Premix Ex Taq™ (Tli RNaseH Plus) (Takara Clontech) on a Light Cycler 480 SYBR Green I master and a Light Cycler 480 apparatus (both from Roche Diagnostics, Indianapolis, IN) with the following primers: human CD206 forward: 5’-GGGTTGCTATCACTCTCTATG-3’; human CD206 reverse: 5’-TTTCTTGTCTGTTGCCGTAGTT-3’; human GAPDH forward: 5’-GAAGGTCGGAGTCAACGGATT-3’; human GAPDH reverse: 5’-TGACGGTGCCATGGAATTTG-3’. *CD206* expression was normalized to *GAPDH* expression.

### Supplementary legends to Tables and Figures

**Figure S1. Relapse-free survival in function of the SPARC expression status in TNBC cancer cells.** Patients with TNBC were divided in two subgroups according to SPARC expression in tumor cells: SPARC^+^ and SPARC^-^.

**Figure S2. Relapse-free survival in function of SPARC status in TAMs.** Patients with TNBC were divided in two subgroups according to SPARC expression in TAMs: SPARC^+^ and SPARC^-^.

**Figure S3. Relapse-free survival according in function of SPARC expression status in endothelial cells.** Patients with TNBC were divided in two subgroups according to SPARC expression in endothelial cells within the tumor microenvironment: SPARC^+^ and SPARC^-^.

**Figure S4. Relapse-free survival according to SPARC expression status in TILs.** Patients with TNBC were divided in two subgroups according to SPARC expression in TILs: SPARC^+^ and SPARC^-^.

**Figure S5. THP1 monocyte differentiation into M2 macrophages**

mRNA expression of the M2 macrophage marker CD206 was quantified by RT-qPCR in THP1 monocytes, M0-, and M2-polarized THP1 macrophages. Data were normalized to *GAPDH* expression level. Results are expressed as mean ± SD (n = 3).

**Figure S6. Expression of *SPARC* and *POSTN* mRNAs in TNBC by single-cell RNA-seq analysis**

**(A) Cell populations.** Thirteen cell populations were identified by single-cell RNA-seq analysis of the previously published GSE118390 dataset (n=6 TNBC samples), according to [9].

**(B) *SPARC* and *POSTN* mRNA expression.** *SPARC* and *POSTN* mRNA relative expressions in each of the 13 cell populations identified by single-cell RNA-seq analysis according to [9]. Individual cell populations were annotated as published in the original single-cell RNA-seq study [9] except for TNBC cells that are labelled Cancer_P1, _P2, _P3, _P4, _P5, _P6.

**Figure S7. Expression of *SPARC* and *POSTN* mRNAs in CAF-S1 clusters in breast cancer by single-cell RNA-seq analysis**

Eight CAF-S1 clusters were identified in the previously published single-cell RNA-seq dataset EGAS00001004030 (n=6 primary breast cancer samples) according to [10]. *SPARC* and *POSTN* mRNA relative expression in the eight CAF-S1 clusters, according to [10], is shown. Individual CAF-S1 clusters were annotated as published in the original single-cell RNA-seq study [10]. According to [10], myCAFs were identified in the following five clusters: ECM-myCAF: associated with extracellular matrix (ECM) remodeling, cell-substrate adhesion, and collagen formation; TGFβ-myCAF: associated with TGFβ signaling pathway and matrisome; Wound-myCAF: associated with assembly of collagen fibrils and wound healing; IFNαβ-myCAF: associated with IFNα/β signaling; Acto-myCAF: associated with the actomyosin complex. iCAF were identified in the following three clusters: Detox-iCAF: associated with the detoxification and inflammatory responses; IL-iCAF: associated with the response to growth factors, TNF signaling, and interleukin (IL) pathway; IFNγ-iCAF: associated with the response to IFNγ and cytokine-mediated signaling pathways.

**Figure S8. Co-localization of SPARC with periostin in TNBC PDX**

PDX B1995 tumor sections were co-incubated with an anti-SPARC polyclonal antibody (15274-1-AP) (red) and an anti-periostin monoclonal antibody (Proteintech) (green). Nuclei were stained with Hoechst 33342 (blue). Top panels: SPARC, periostin, and merge. Bottom panels: higher magnification of the boxed areas Arrows indicate SPARC and periostin co-localization. Scale bar, 10 μm.

**Figure S9. Immunodepletion of SPARC secreted in the conditioned medium from HMFs.** SPARC was immunodepleted or not in conditioned medium (CM) from HMFs (CM HMF) by immunoprecipitation. SPARC immunodepletion in CM HMF (CM HMF IP SPARC) was confirmed by western blot analysis with an anti-SPARC monoclonal antibody (clone AON-5031).

**Table S1. Univariate and multivariate logistic regression analyses to identify prognostic factors of overall survival (OS) in TNBC.**

**Table S2. Clinicopathological characteristics in function of SPARC expression (SPARC^+^ and SPARC^-^) in CAFs.**

