## Supplementary figures and images for "SPARC in cancer-associated fibroblasts is an independent poor prognostic factor in non-metastatic triple-negative breast cancer and exhibits pro-tumor activity"

### supplemental figure 1

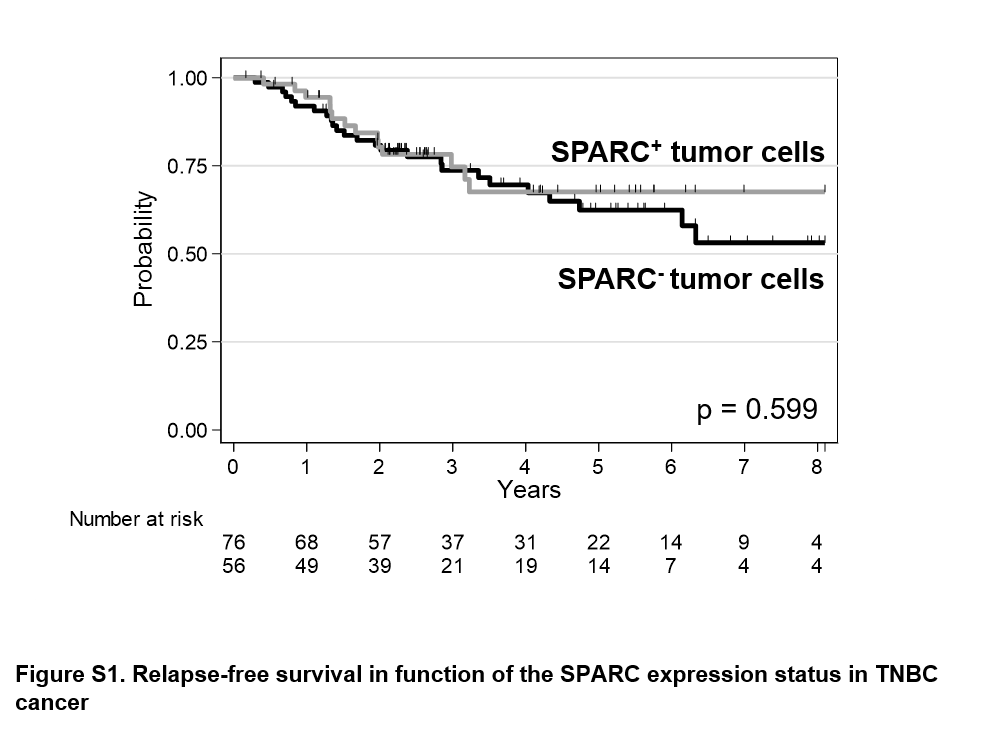

### supplemental figure 2

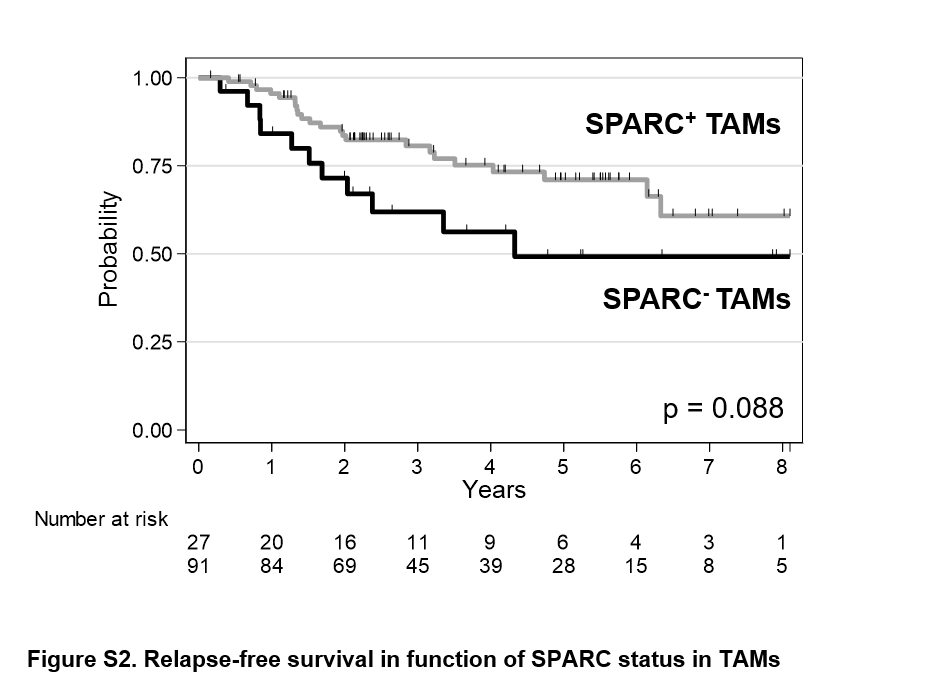

### supplemental figure 3

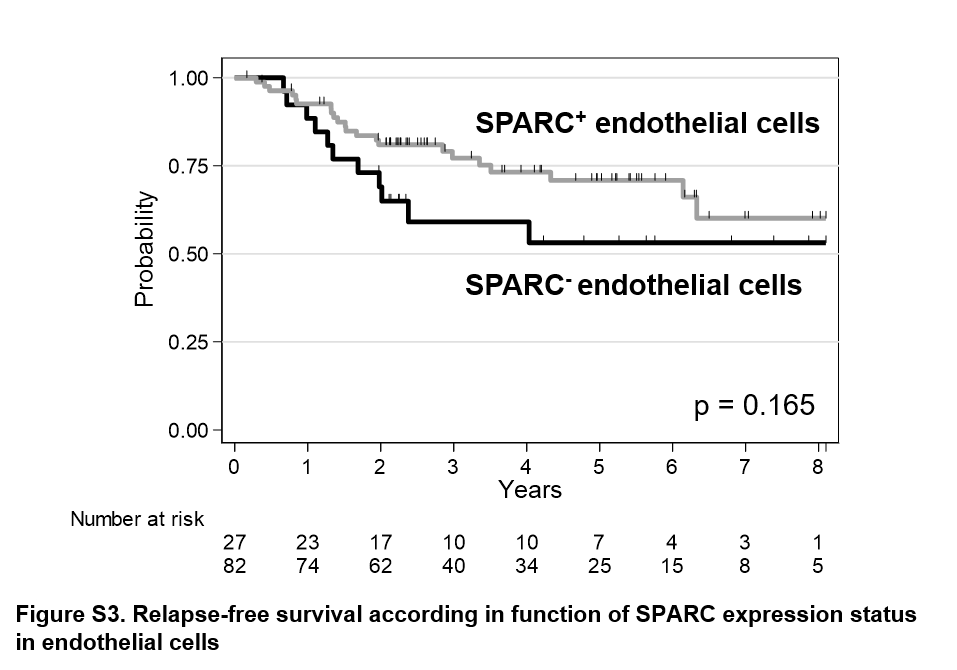

### supplemental figure 4

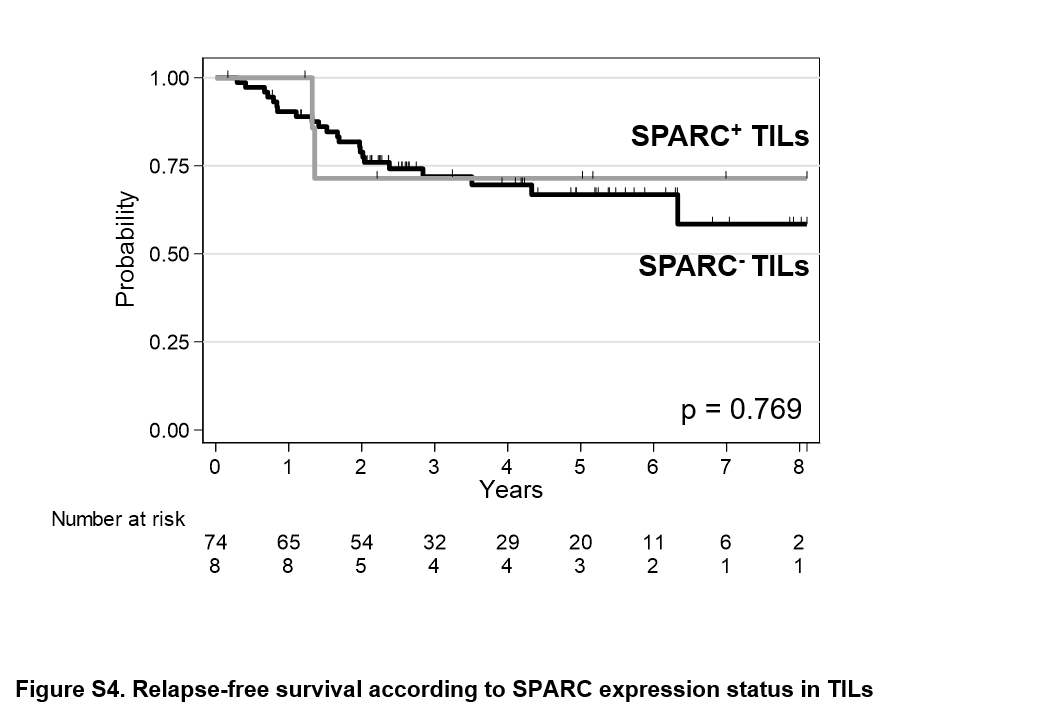

### supplemental figure 5

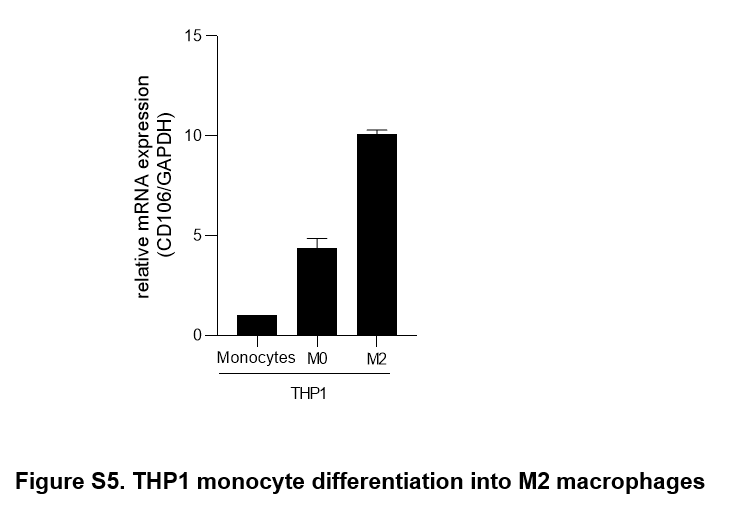

### supplemental figure 6

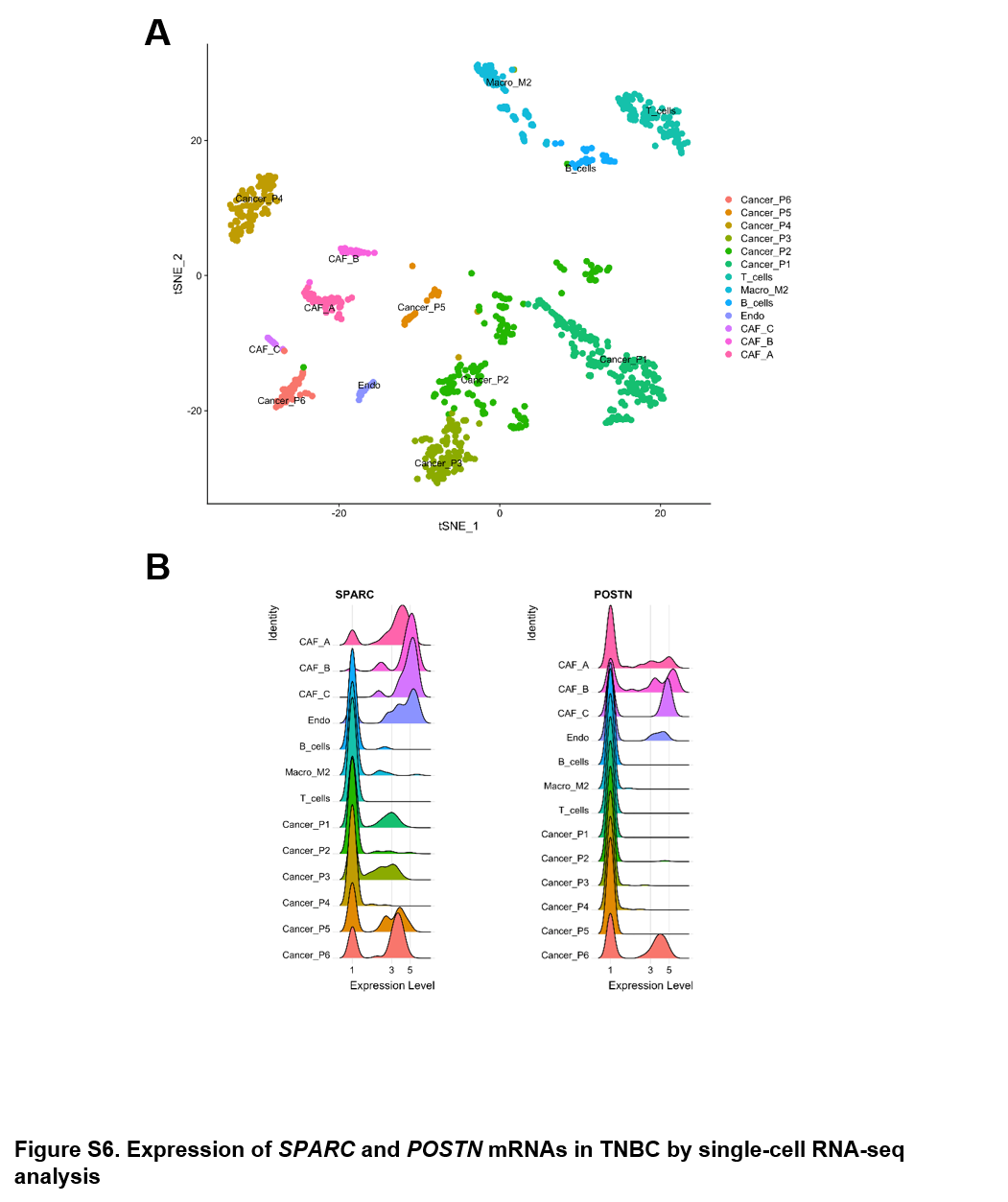

### supplemental figure 7

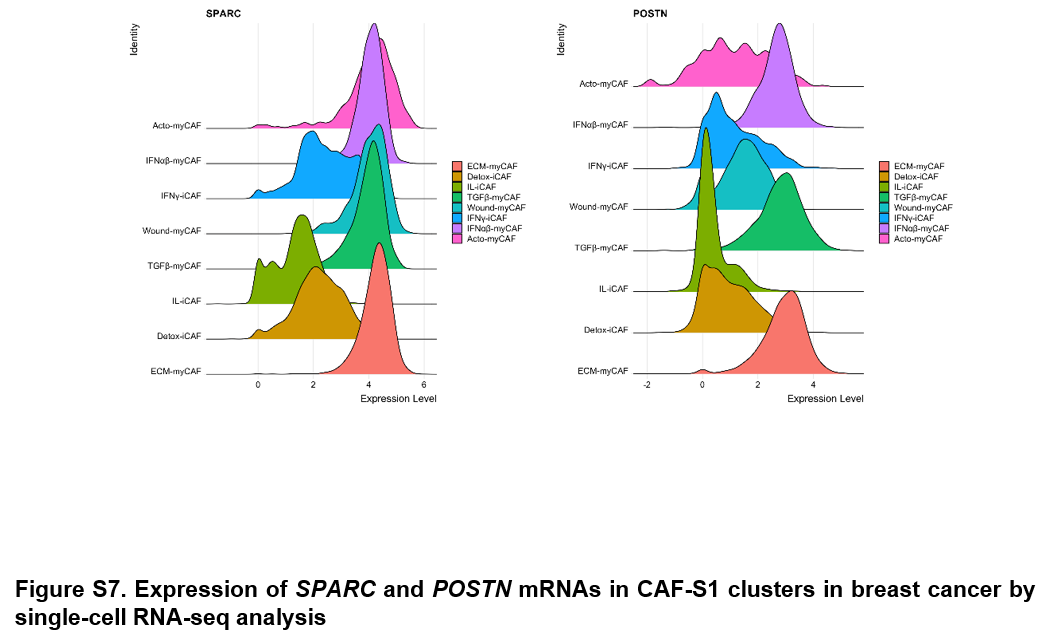

### supplemental figure 8

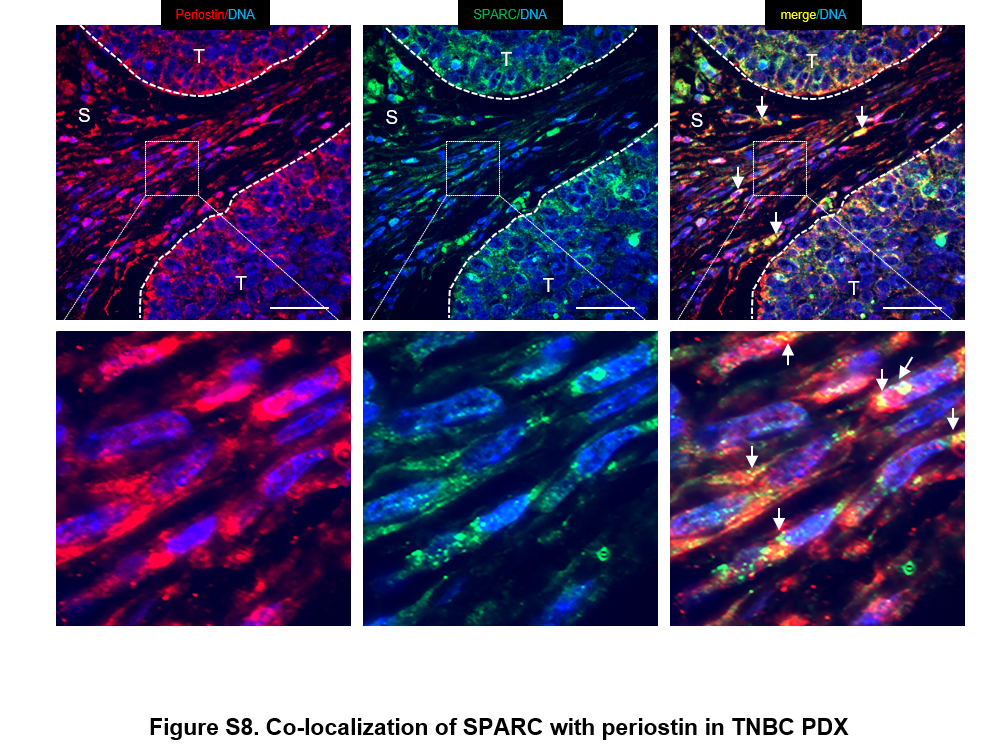

### supplemental figure 9

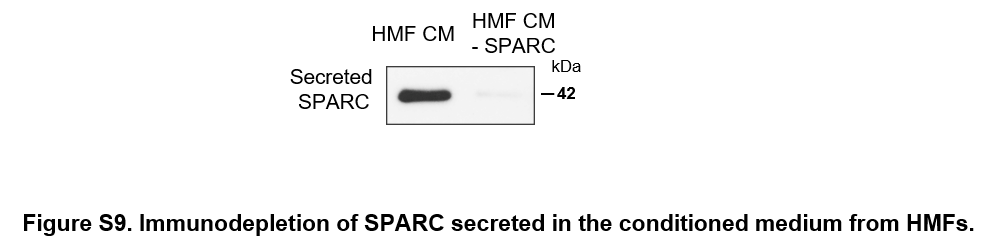
