## supplemental table S1 for "SPARC in cancer-associated fibroblasts is an independent poor prognostic factor in non-metastatic triple-negative breast cancer and exhibits pro-tumor activity"

| Clinical and tumor characteristics | Univariate analysis HR 95% CI | Multivariate analysis HR 95% CI |
|---|---|---|
| **Age** | N=148 | |
| < 55 years | 1 | |
| ≥ 55 years | 2.21 [1.03-4.75] | |
| | ***P* = 0.027** | |
| **Tumor size** | N=148 | |
| T1 | 1 | 1 |
| T2 | 3.04 [1.17-7.93] | 1.99 [0.74-5.30] |
| T3/T4 | 6.73 [2.34-19.38] | 3.66 [1.22-11.0] |
| | ***P* < 0.001** | ***P* = 0.050** |
| **Nodal status** | N=148 | |
| N- | 1 | 1 |
| N+ | 2.47 [1.36-4.47] | 2.38 [1.25-4.55] |
| | ***P* = 0.002** | ***P* = 0.008** |
| **Histological grade (SBR)** | N=146 | |
| 1-2 | 1 | |
| 3 | 1.07 [0.45-2.53] | |
| | *P* = 0.883 | |
| **Histology** | N=145 | |
| Ductal | 1 | |
| Lobular | 0.70 [0.22-2.29] | |
| Other | 1.17 [0.42-3.29] | |
| | *P* = 0.777 | |
| **Adjuvant chemotherapy** | N=148 | |
| No | 1 | 1 |
| Yes | 0.36 [0.20-0.64] | 0.33 [0.18-0.60] |
| | ***P* = 0.006** | ***P* < 0.001** |

| | |
|---|---|
| **Basal-like phenotype**<br>Yes<br>No | N=147<br>1<br>1.16 [0.64-2.08]<br>*P* = 0.624 |
| **SPARC expression in tumor cells**<br>Negative<br>Positive | N=132<br>1<br>0.75 [0.40-1.41]<br>*P* = 0.363 |
| **SPARC expression in CAFs**<br>Negative<br>Positive | N=126<br>1<br>1.92 [0.59-6.23]<br>*P* = 0.235 |
| **SPARC expression in TAMs**<br>Negative<br>Positive | N=118<br>1<br>0.59 [0.29-1.16]<br>***P* = 0.139** |
| **SPARC expression in endothelial cells**<br>Negative<br>Positive | N=109<br>1<br>0.73 [0.35-1.51]<br>*P* = 0.401 |
| **SPARC expression in TILs**<br>Negative<br>Positive | N=82<br>1<br>1.10 [0.32-3.75]<br>*P* = 0.882 |
| **TIL density**<br>[0-1]<br>>1 | N=142<br>1<br>0.94 [0.50-1.75]<br>*P* = 0.839 |
| **PDL-1 expression in tumor cells**<br>< 1%<br>≥ 1% | N=136<br>1<br>0.91 [0.49-1.70]<br>*P* = 0.770 |

| | |
|---|---|
| **PDL-1 expression in TILs** | N=134 |
| 0 | 1 |
| ]0-50[ | 1.90 [0.66-5.45] |
| ≥ 50 | 1.10 [0.33-3.66] |
| | *P* = 0.238 |
| **PD1 expression in TILs** | N=140 |
| 0 | 1 |
| ]0-50[ | 0.94 [0.40-2.25] |
| ≥ 50 | 0.73 [0.22-2.38] |
| | *P* = 0.832 |
| **Fibrosis** | N=137 |
| ≤ 50% | 1 |
| > 50% | 1.10 [0.60-2.02] |
| | *P* = 0.746 |
| **TAMs (inflammation)** | N=143 |
| 0/1 | 1 |
| 2 | 1.11[0.51-2.43] |
| 3 | 0.67 [0.31-1.45] |
| | *P* = 0.285 |

**Table S1. Univariate and multivariate logistic regression analyses to identify prognostic factors of overall survival (OS) in TNBC**

SBR: Scarff-Bloom-Richardson; CAFs: cancer-associated fibroblasts; TAMs: tumor-associated macrophages; TILs: tumor-infiltrating lymphocytes
HR = hazard ratio; CI = confidence interval; p values in bold, statistically significant.
