## supplemental table S2 for "SPARC in cancer-associated fibroblasts is an independent poor prognostic factor in non-metastatic triple-negative breast cancer and exhibits pro-tumor activity"

| Clinical and tumor characteristics | SPARC expression in CAFs Negative (N=15) | | SPARC expression in CAFs Positive (N=111) | | *P value* |
|---|---|---|---|---|---|
| **Age** | | | | | 0.018 |
| < 55 years | 1 | (6.7%) | 43 | (38.7%) | |
| ≥ 55 years | 14 | (93.3%) | 68 | (61.3%) | |
| **Tumor size** | | | | | 0.334 |
| T1 | 6 | (40.0%) | 36 | (32.4%) | |
| T2 | 9 | (60.0%) | 59 | (53.2%) | |
| T3/T4 | 0 | | 16 | (14.4%) | |
| **Nodal status** | | | | | 0.863 |
| N- | 9 | (60.0%) | 64 | (57.7%) | |
| N+ | 6 | (40.0%) | 47 | (42.3%) | |
| **Histological grade (SBR)** | | | | | 0.130 |
| 1-2 | 3 | (20.0%) | 8 | (7.3%) | |
| 3 | 12 | (80.0%) | 101 | (92.7%) | |
| **Histology** | | | | | 0.080 |
| Ductal | 11 | (73.3%) | 95 | (88.0%) | |
| Lobular | 3 | (20.0%) | 5 | (4.6%) | |
| Other | 1 | (6.7%) | 8 | (7.4%) | |
| **Adjuvant chemotherapy** | | | | | 0.186 |
| No | 7 | (46.7%) | 33 | (29.7%) | |
| Yes | 8 | (53.3%) | 78 | (70.3%) | |

| | | | |
|---|---|---|---|
| **Basal-like phenotype** | | | 0.678 |
| ≤ 10% | 9 (60.0%) | 72 (65.4%) | |
| Basal | 6 (40.0%) | 38 (34.6%) | |
| **SPARC expression in tumor cells** | | | 0.603 |
| Negative | 8 (53.3%) | 67 (60.4%) | |
| Positive | 7 (46.7%) | 44 (39.6%) | |
| **SPARC expression in TAMs** | | | **0.007** |
| Negative | 7 (58.3%) | 20 (19.4%) | |
| Positive | 5 (41.7%) | 83 (80.6%) | |
| **SPARC expression in endothelial cells** | | | **0.026** |
| Negative | 7 (50.0%) | 20 (22.0%) | |
| Positive | 7 (50.0%) | 71 (78.0%) | |
| **SPARC expression in TILs** | | | 1.000 |
| Negative | 8 (100.0%) | 65 (89.0%) | |
| Positive | 0 | 8 (11.0%) | |
| **TIL density** | | | 0.127 |
| [0-1] | 6 (42.9%) | 26 (23.9%) | |
| > 1 | 8 (57.1%) | 83 (76.1%) | |
| **PDL-1 expression in tumor cells** | | | 0.109 |
| < 1% | 7 (53.9%) | 31 (28.4%) | |
| ≥ 1% | 6 (46.1%) | 78 (71.6%) | |

| | | | | | |
|---|---|---|---|---|---|
| **PDL-1 expression in TILs** | | | | | **0.049** |
| 0 | 4 | (30.8%) | 10 | (9.2%) | |
| ]0-50[ | 7 | (53.8%) | 61 | (56.0%) | |
| ≥ 50 | 2 | (15.4%) | 38 | (34.8%) | |
| **PD1 expression in TILs** | | | | | 0.415 |
| 0 | 2 | (15.4%) | 11 | (10.2%) | |
| ]0-50[ | 8 | (61.5%) | 83 | (76.9%) | |
| ≥ 50 | 3 | (23.1%) | 14 | (12.9%) | |
| **Fibrosis** | | | | | **0.028** |
| ≤ 50% | 3 | (20.0%) | 56 | (51.4%) | |
| > 50% | 12 | (80.0%) | 53 | (48.6%) | |
| **TAMs (inflammation)** | | | | | 0.349 |
| 0/1 | 4 | (28.6%) | 15 | (13.8%) | |
| 2 | 3 | (21.4%) | 32 | (29.4%) | |
| 3 | 7 | (50.0%) | 62 | (56.9%) | |

**Table S2**. **Clinicopathological characteristics in function of SPARC expression ($SPARC^+$ and $SPARC^-$) in CAFs**

SBR: Scarff-Bloom-Richardson; CAFs: cancer-associated fibroblasts; TAMs: tumor-associated macrophages; TILs: tumor-infiltrating lymphocytes.
